# Genome Mining and Discovery of Imiditides, a Novel Family of RiPPs with a Class-defining Aspartimide Modification

**DOI:** 10.1101/2023.04.07.536058

**Authors:** Li Cao, Truc Do, Angela D. Zhu, Nathan Alam, A. James Link

## Abstract

Ribosomally synthesized and post-translationally modified peptides (RiPPs) are a fascinating class of natural products of ribosomal origins. In the past decade, various sophisticated machine learning-based software packages have been established to discover novel RiPPs that do not resemble the known families. Instead, we argue that tailoring enzymes that cluster with various RiPP families can serve as effective bioinformatic seeds for novel RiPP discovery. Leveraging that *O*-methyltransferases homologous to protein isoaspartyl methyltransferases (PIMTs) are associated with lasso peptide, graspetide, and lanthipeptide biosynthetic gene clusters (BGCs), we utilized the C-terminal motif unique to RiPP-associated *O*-methyltransferases as the search query to discover a novel family of RiPPs, imiditides. Our genome-mining algorithm reveals a total of 670 imiditide BGCs, widely distributed in Gram-positive bacterial genomes. In addition, we demonstrate the heterologous production of the founding member of the imiditide family, mNmaA^M^, encoded in the genome of *Nonomuraea maritima*. In contrast to other RiPP associated PIMTs that recognize constrained peptides as substrates, the PIMT homolog in mNmaA^M^ BGC, NmaM, methylates a specific Asp residue on the linear precursor peptide, NmaA. The methyl ester is then turned into an aspartimide spontaneously. The aspartimide moiety formed is unusually stable, leading to the accumulation of the aspartimidylated product *in vivo*. The substrate specificity is achieved by extensive charge-charge interactions between the precursor NmaA and the modifying enzyme NmaM suggested by both experimental validations as well as an AlphaFold model prediction. Our study suggests that PIMT-mediated aspartimide formation is an underappreciated backbone modification strategy in RiPP biosynthesis, compared to the well-studied backbone rigidification chemistries, such as thiazol(in)e and oxazol(in)e formations. Additionally, our findings suggest that aspartimide formation in Gram-positive bacterial proteomes are not limited to spontaneous protein aging and degradation.

**TOC Figure:** 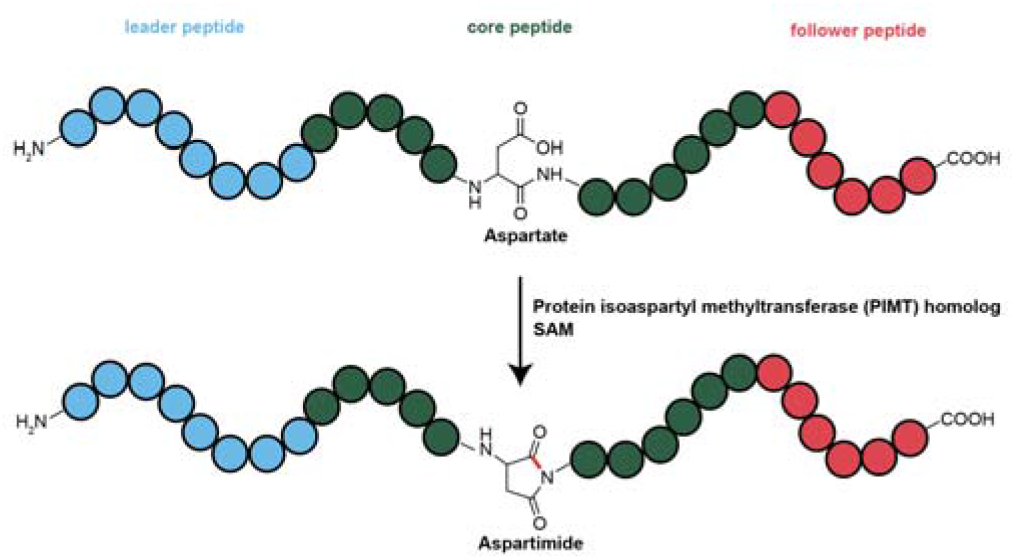

---

Ribosomally synthesized and post-translationally modified natural products (RiPPs) are a diverse class of natural products of ribosomal origin.^1–3^ RiPP biosynthesis begins with a ribosomally synthesized precursor peptide, whose N-terminal leader peptide recruits the tailoring enzymes to post-translationally modify the C-terminal core sequences.^1,2^ As the precursor sequences of RiPPs are gene-encoded, genome-mining approaches have proven to be an exceptionally well-suited strategy for RiPP discoveries.^2,4–9^ In fact, genome mining for RiPPs has advanced to the point where identification of well-known RiPPs such as lanthipeptides, lasso peptides, and thiopeptides can be readily identified from genome sequences using tools like antiSMASH.^4,10^ Improvements in genome annotation have also led to the assignment of many genes involved in RiPP biosynthesis in the genome sequences deposited in GenBank.

The next frontier in genome mining for RiPPs has focused on the discovery of entirely new classes of RiPPs that do not resemble the known families. Sophisticated machine learning-based software packages have been developed for this task.^11,12^ Previous work by our group and the van der Donk group reported homologs of protein isoaspartyl methyltransferases (PIMTs) are associated with biosynthetic gene clusters of several RiPP families, including lanthipeptides,^13,14^ lasso peptides,^15,16^ and graspetides.^17,18^ We reasoned that these shared auxiliary enzymes associated with known RiPPs could also be used as a bioinformatic handle to identify novel RiPP BGCs. In particular, the PIMT homologs previously reported recognize constrained RiPPs as substrates to install the aspartimide posttranslational modification. Therefore, we investigated whether these PIMT homologs can directly modify linear precursors and become a class-defining enzyme for an uncharacterized family of RiPPs.

Here, we report the discovery and genome-mining of a novel family of RiPPs in which aspartimide formation is the class-defining modification, for which we propose the name imiditide. The results of our domain-centric genome mining reveal 670 imiditide BGCs, all of which are in Gram-positive bacterial genomes. Representative genera that encode imiditides include *Streptomyces, Actinomadura*, and *Nonomuraea*. In addition, we report the heterologous production of an imiditide from *Nonomuraea maritima* in *E. coli*. The PIMT homolog in the BGC, NmaM, directly methylates a specific aspartate residue on the linear precursor, NmaA. The aspartimide is then spontaneously formed. The aspartimide in the resulting imiditide exhibits some stability, but can be hydrolyzed regioselectively to incorporate the isoaspartate residue into the polypeptide. Our experimental evidence along with the AlphaFold model of the NmaA-NmaM complex suggest that the specificity of the modification is achieved through charge-charge interactions. Our findings demonstrate the feasibility of discovering novel classes of RiPPs by using shared tailoring enzymes across different RiPP families as bioinformatic seeds. Furthermore, we illustrate that PIMT-mediated aspartimide formation is an underappreciated backbone modification strategy in RiPP biosynthesis, compared to the well-studied backbone rigidification chemistries, such as thiazol(in)e and oxazol(in)e formations.^19,20^

## Results

### Identification of Biosynthetic Gene Clusters for a Putative Novel RiPP Family

Our laboratory has previously characterized several RiPPs with aspartimide post-translational modifications, including lasso peptides cellulonodin-2 and lihuanodin,^15^ as well as graspetides fuscimiditide and amycolimiditide.^17,21^ In their biosynthesis, a dedicated *O*-methyltransferase related to protein isoaspartyl methyltransferases (PIMT) installs the aspartimide on the matured lasso peptides or graspetides as the last step of the reaction series. Since PIMT homologs cluster with various families of RiPPs (Figure 1a), we reason that these PIMT homologs can serve as a bioinformatic handle to unveil uncharacterized families of RiPPs.

**Figure 1:**
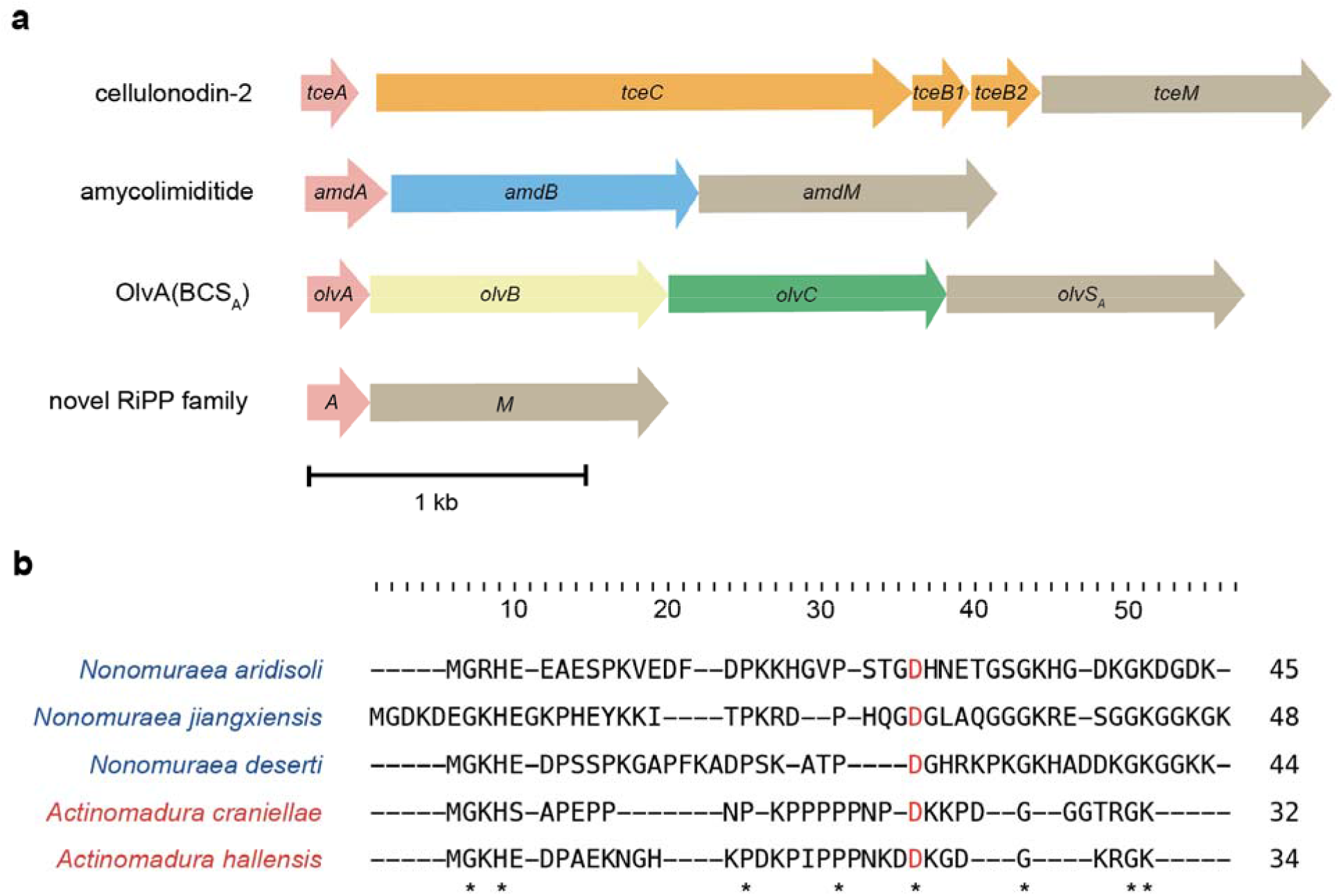
Identification of a Novel RiPP Family. **a)** Biosynthetic gene clusters containing PIMT homologs. Red, precursor peptides; brown, *O*-methyltransferases that are PIMT homologs. In the cellulonodin-2 BGC, *tceB1* and *tceB2* code for a bipartite cysteine peptidase, while *tceC* encodes the lasso cyclase (orange). In the amycolimiditide BGC, *amdB* encodes an ester-installing ATP-grasp enzyme. In the OlvA(BCS_A_) BGC, *olvB* encodes a dehydratase, and *olvC* codes a cyclase. Identified here first, BGCs corresponding to the novel RiPP family only harbor genes encoding a precursor and an *O*-methyltransferase. **b)** Sequence alignments of 5 putative precursor sequences in the novel RiPP family. *Nonomuraea* strains are colored blue and *Actinomadura* strains are colored red. The putative site of modification (Asp) is colored red. Conserved residues include Gly, His, Pro, and Lys, which are marked with asterisks.

We first constructed sequence similarity network (SSN) of ThfM (Figure S1), the *O*-methyltransferase responsible for installing the aspartimide in graspetide fuscimiditide (Figure 1a).^17^ A total of 2000 sequences were queried to construct the SSN. ThfM surprisingly does not cluster with any other sequence. On the other hand, AmdM, the methyltransferase installing an aspartimide in tetracyclic graspetide amycolimiditide, clusters with methyltransferases from phylogenetically relevant strains that are also near other graspetide BGCs. Similarly, OlvS_A_, the PIMT homolog responsible for the isoaspartate incorporation in OlvA(BCS_A_) through the aspartimide formation, clusters with other methyltransferases near lanthipeptide BGCs. TceM and LihM are not a part of this sequence similarity network, showing they are more distant from ThfM. Additionally, we noticed a cluster that contain a large number of PIMT homologs from *Nonomuraea* and *Actinomadura* strains. These PIMT homologs appear to be isolated in the genome context at first glance. However, we observed an unannotated open reading frame (ORF) of 30-75 amino acoids that neighbors with the PIMT homolog. To investigate their potential as class-defining enzymes for a novel family of RiPPs, we collected a set of 50 biosynthetic gene clusters (BGCs) by performing a BLASTP search on an *O*-methyltransferase from *Nonomuraea jiangxiensis* found in the ThfM SSN (Table S1). The putative precursor sequences in these BGCs contain the Asp residue required to be recognized by the PIMT homolog and are also rich in Gly, Lys and Pro residues (Figure 1b). Apart from genes encoding the putative precursor and the PIMT homolog, there are no other conserved genes in these BGCs. Therefore, we hypothesize that the PIMT homolog in these BGCs can be the class-defining enzymes for a novel family of RiPPs, where the *O*-methyltransferase directly acts on the linear precursor to install a posttranslational modification.

### Genome Mining of the Biosynthetic Gene Clusters (BGCs) for the Putative Novel RiPP Family

*O*-methyltransferases near RiPP BGCs are often annotated as methyltransferase domain-containing protein on NCBI and consist of an N-terminal region that is homologous to PIMT (∼220 aa) and a C-terminal domain of 110-180 aa (Figure S2). In our previous study on cellulonodin-2 and lihuanodin, a conserved WXXXGXP motif was identified in the C-terminal domain of PIMT homologues near lasso peptide BGCs. A disruption of this motif in the TceM and LihM methyltransferases has a deleterious effect on aspartimidylation of pre-cellulonodin-2 and pre-lihuanodin, showing that the C-terminal domain plays an important role in substrate recognition.^15^ Additionally, the length of the C-terminal domain of these methyltransferases varies based on the type of RiPP families with which they cluster, and the sequences of methyltransferases associated with the same family show much more resemblance (Figure S2).

To probe if the 50 PIMT homologs residing within a putative RiPP BGC (Table S1) have a similarly conserved sequence motif, we first subjected these sequences to Multiple EM for Motif Elicitation (MEME).^22^ MEME analysis revealed a conserved 41 aa sequence motif within the C-terminal domain of the PIMT homologs (Figure 2a). A limited BLASTP search with this motif then revealed that it is a unique signature of methyltransferases encoded within minimal BGCs for the novel RiPP family. As such, we reasoned that we could use this conserved motif as a bioinformatic handle for genome mining of this new class of PIMT-associated RiPPs.

**Figure 2:**
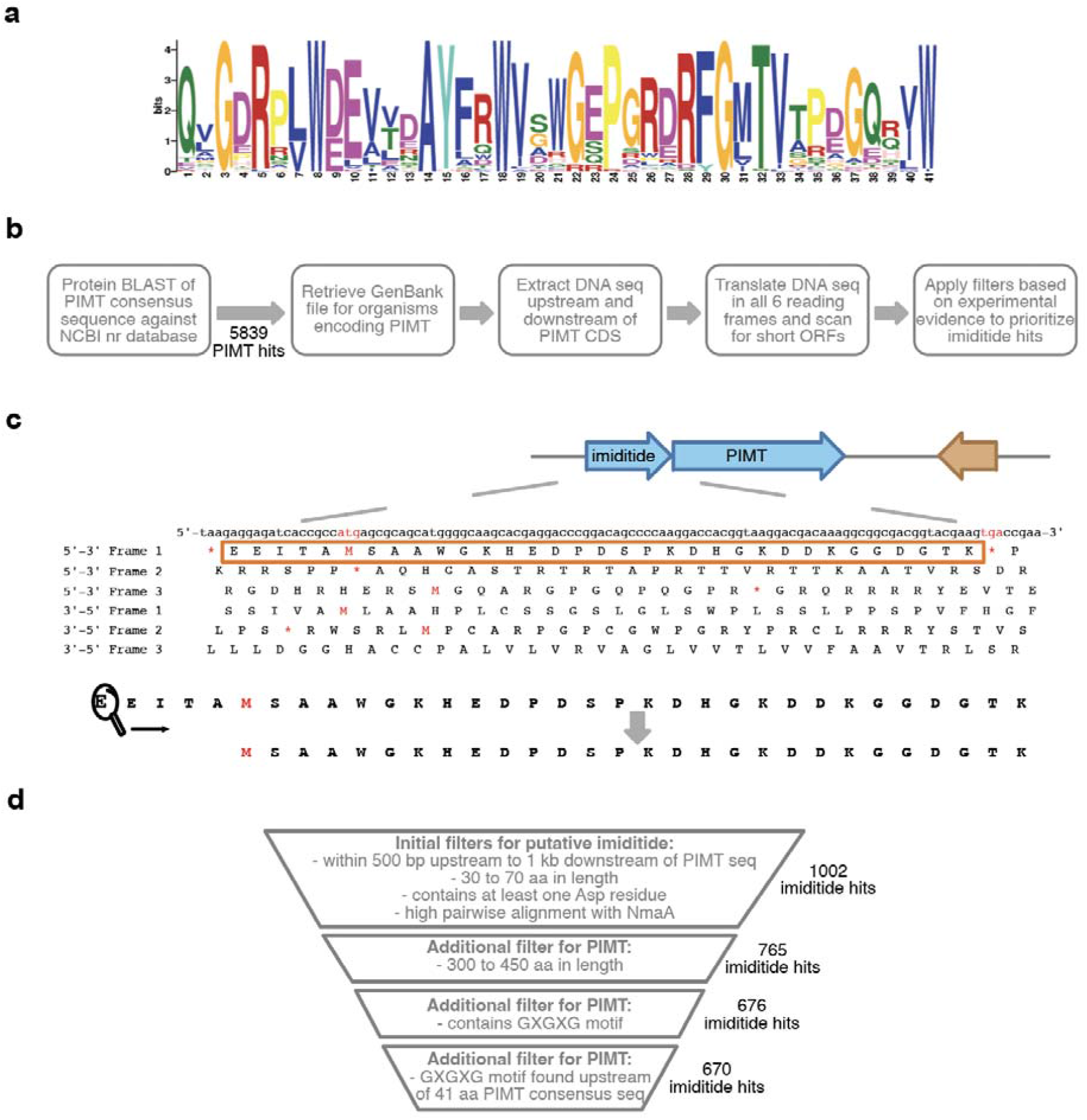
Genome Mining of Imiditide Biosynthetic Gene Clusters. **a)** Conversed Sequence Logo of a conserved 41 aa C-terminal motif of 50 methyltransferases associated with putative imiditide BGCs. A complete list of 50 methyltransferases is shown in Table S1. **b)** Overall strategy for the genome mining of imiditide BGCs. Blasting the 41 aa conserved sequences against the NCBI protein database returns a total of 5839 hits. Then, we search short open reading frames that contain putative precursor sequences near these PIMT homologs. **c)** An example workflow to identify the putative precursor sequence near a PIMT homolog. All 6 frames are translated, and putative precursor sequences are searched within 500 upstream or 1000 downstream of the PIMT homologs. Only the ORFs that contains at least one Asp residue and have a size between 30 to 75 aas will suffice as putative precursors. **d)** Applying additional filters based on other PIMT-associated RiPP clusters further eliminate potential false positive hits, resulting a total of 670 identified imiditide BGCs.

As the first step in our genome mining pipeline, we comprehensively identified all PIMT homologs harboring the unique 41 aa motif by BLASTP search of the motif against the entire non-redundant NCBI protein database. Using an e-value cutoff of 1000, 5839 hits were obtained. For each of the PIMT homologs identified, we searched for an associated RiPP coding sequence in the vicinity by scanning for short open reading frames. Based on previous work, we initially filtered for ORFs encoding a 30-75 aa product within 500 bp upstream or 1000 bp downstream of the PIMT homolog and having at least one Asp residue in the protein sequence (Figure 2b).^6^ Given the operon nature of BGCs, the short ORF and PIMT homolog coding sequence were also required to be arranged in parallel on the same DNA strand. When multiple short ORFs were identified within the defined windows of a PIMT homolog, only the ORF that has the highest alignment score with the putative precursor sequence from *Nonomuraea maritima* (Figure 1b) was considered, as the cluster was experimentally verified in the later sections. This analysis generated 1002 potential PIMT-associated RiPPs, hereafter referred to as imiditides (hit rate of 17.2%). To further reduce potential false positive hits, we applied two additional filters to eliminate predicted imiditide BGCs that lack a close PIMT homolog. Based on observation of experimentally characterized RiPP-associated methyltransferases, we limited the length of PIMT homologs to 300-450 aa inclusive (Figure S2). Additionally, we required that the PIMT homologs harbor a GXGXG motif upstream of the 41 aa motif used in our initial protein BLAST search; the GXGXG motif has been shown to bind SAM in human PIMT (Figure S3). After applying these additional filters, we identified 670 putative imiditide BGCs containing a PIMT homolog, all in Gram-positive bacteria. The most represented genus from which these BGCs are found include *Streptomyces, Actinomadura, Nonomuraea*, and *Nocardiopsis* (Figure S4). Notably, all RiPP-associated PIMT homologs are also encoded in Gram-positive bacteria.

### Heterologous Expression of Imiditide from *Nonomuraea maritima*

To validate these putative BGCs indeed encode RiPPs, we attempted to heterologously express a putative imiditide BGC from *Nonomuraea maritima* in *E. coli*. In the native BGC, the ORF that encodes the precursor, *nmaA*, is immediately upstream of the gene encodes the methyltransferase, *nmaM* (Figure 3a, c). The putative precursor sequence NmaA is predicted to be 46 aa long, and the methyltransferase NmaM has a size of 373 aa. To enhance the expression level of NmaA, we refactored the cluster by adapting a co-expression system. His_6_-SUMO-NmaA was placed under an IPTG-inducible T5 promoter, while untagged NmaM was put under an IPTG-inducible T7 promoter on a separate plasmid (Figure 3b). Then, we coexpressed these two plasmids using *E. coli* BL21(DE3) Δ*slyD* strain at room temperature for 20 hours (Figure 3b). Products are purified using nickel affinity chromatography under the urea denaturing condition. Chromatogramming the purified fraction on LC-MS revealed the formation of two additional species, with one carrying an addition of 14 Da and the other one bearing a loss of 18 Da (Figure 3d). These species were absent when we expressed His_6_-SUMO-NmaA alone, indicating that the formation of these species was due to the action of NmaM (Figure 3d). Additionally, the SDS-PAGE gel of the purified protein fractions revealed that modified His_6_-SUMO-NmaA pulled down untagged NmaM even in the presence of 8M urea, indicating an exceptionally strong binding between the two proteins (Figure 3e). Since NmaM is predicted to be a PIMT homolog, we hypothesize that NmaM methylates a specific Asp side chain, followed by nucleophilic attack of the adjacent amide to form an aspartimide (Figure 3f). The masses of these two new species agreed with this hypothesis, as one was putatively methylated (+14 Da) and the other one was putatively aspartimidylated (−18 Da) (Figure 3d). Additionally, to confirm that the SUMO tag did not interfere with the post-translational modification, we coexpressed His_6_-NmaA with NmaM, and also observed the emergence of the putative methylated and aspartimidylated species (Figure S5). Since the SUMO tag greatly improved the expression level of NmaA, we used His_6_-SUMO-NmaA for all subsequent experiments.

**Figure 3:**
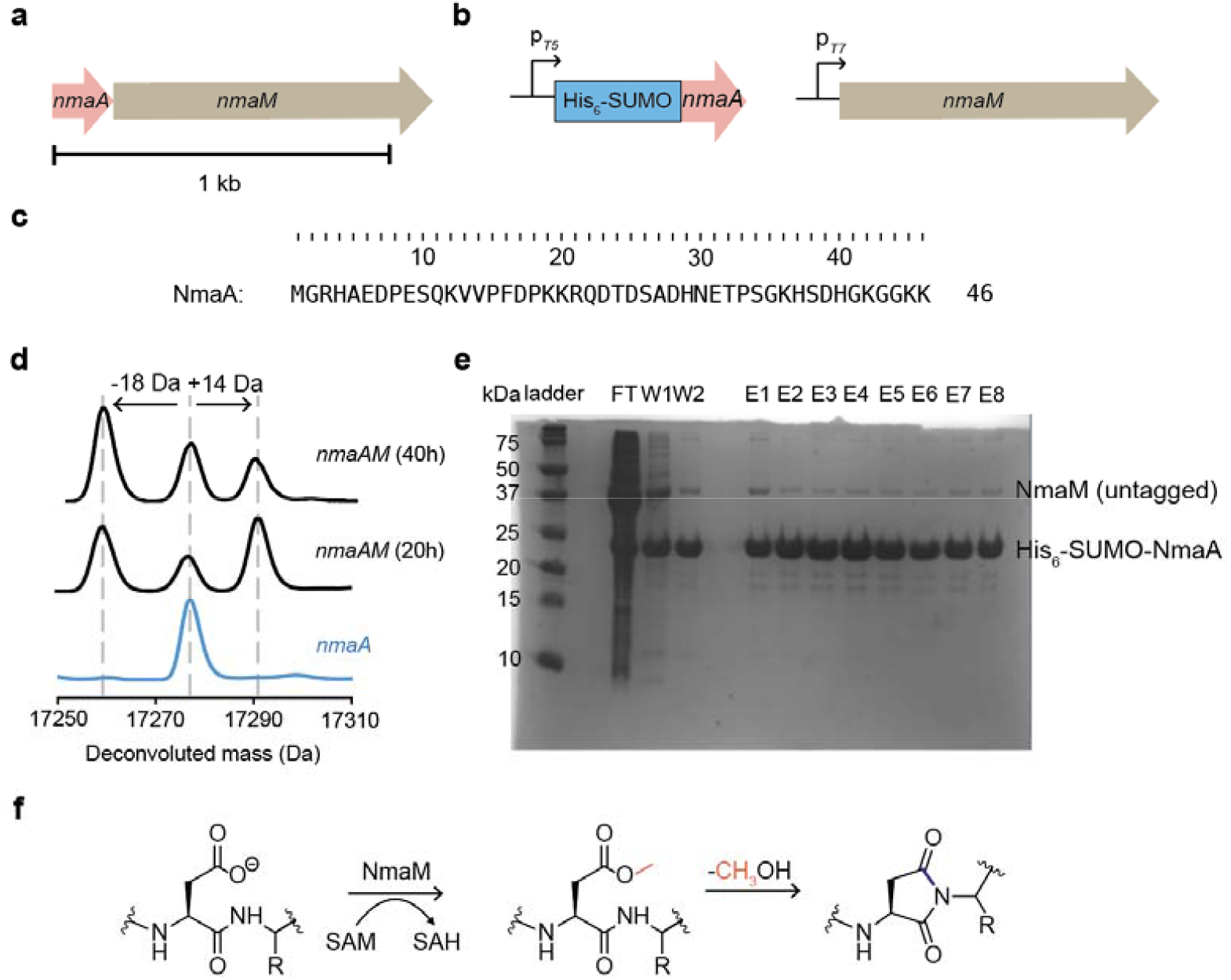
Hetereologous Expression of mNmaA^M^. **a)** Native gene cluster of a putative imiditide from *Nonomuraea maritima. nmaA* encodes the putative precursor sequence (sequence shown in part c), and *nmaM* encodes an *O*-methyltransferase that is a PIMT homolog. **b)** Refactored gene clusters for mNmaA^M^. An IPTG inducible T5 promoter was introduced upstream of a N-terminally His_6_-SUMO tagged precursor peptide gene. An IPTG-inducible T7 promoter was used to drive the untagged methyltransferase (NmaM) expression. **c)** Amino acid sequence of the putative precursor NmaA. **d)** NmaM was responsible for the formation of the putative methylated and aspartimidylated species. The heterologous expression of His_6_-SUMO-NmaA alone returned a species with a mass of 17277.1 Da (expected mass: 17277.1 Da). A coexpression of His_6_-SUMO-NmaA and NmaM resulted in additional putative methylated (+14Da) and aspartimidylated species (−18Da). The putative aspartimidylated species accumulated over time, showing it was the intended product *in vivo* (also shown in Figure S6). **e)** The SDS-PAGE gel showed mNmaA^M^ pulled down untagged NmaM even in the presence of 8M urea, indicating that they bind very tightly. **f)** Proposed reaction pathway for mNmaA^M^. NmaM methylates a specific aspartate residue using SAM. The intramolecular aspartimide is then spontaneously formed.

To determine the kinetics of the reaction, we conducted co-expressions of His_6_-SUMO-NmaA and NmaM at additional various time points (1h, 2h, 3h, 4h and 40h). We observed a faster accumulation of the putative methylated species than the putative aspartimidylated species at shorter time points (Figure S6). As mentioned earlier, after 20h of expression, we detected a ratio of approximately 2:1:2 among the putatively aspartimylated, unmodified and methylated NmaA (Figure 3c). With longer expression time (40h), the ratio shifted to approximately 3:2:1 among putatively aspartimidylated, unmodified and methylated species (Figure 3d). Since the putatively aspartimidylated species accumulated when longer expression time was allowed, we considered this species to be the intended product *in vivo*. We named this putative imiditide mNmaA^M^ following the nomenclature used in our study on amycolimiditide, where m stands for modified, and superscripted M stands for the methyltransferase.^21^ Our observation indicated that the rate of methylation was faster than the rate of aspartimidylation for the biosynthesis of mNmaA^M^ (Figure S6).

Since NmaM is homologous to PIMT, we wonder whether Pcm, the housekeeping PIMT in *E. coli*, could alter the intended product *in vivo*. To investigate this, we constructed *E. coli* BL21(DE3) Δ*pcm*Δ*slyD* strain, and expressed mNmaA^M^ using this strain. The product distribution remained the same using either *E. coli* BL21(DE3) Δ*pcm*Δ*slyD* or *E. coli* BL21(DE3) Δ*slyD*, suggesting that Pcm was not involved in the process (Figure S7). We used *E. coli* BL21(DE3) Δ*slyD* for all subsequent experiments.

To determine the effect of the stoichiometry between NmaA and NmaM on the biosynthesis of mNmaA^M^, we constructed a plasmid where the precursor protein NmaA was placed under an IPTG-inducible T5 promoter and NmaM was placed under the constitutive promoter native to the biosynthetic enzymes needed for lasso peptide microcin J25 biosynthesis.^15,23,24^ Expressing NmaM constitutively resulted in a significant decrease in the protein expression level, which led to only a small fraction of NmaA being modified (Figure S8). It also suggests that NmaM has a low turnover number of NmaA, possibly because modified NmaA species bind to NmaM tightly (Figure 3e).

### mNmaA^M^ Contains an Aspartimide Moiety

Based on our previous studies on aspartimide-containing lasso peptides and graspetides, we hypothesize that mNmaA^M^ also contains an aspartimide moiety (Figure 1a, 3d). Aspartimides are often considered as a reaction intermediate catalyzed by canonical PIMTs.^25^ However, heterologous expression of mNmaA^M^ shows an accumulation of the putative aspartimidylated product *in vivo*, illustrating the relative stability of the moiety in this linear peptide. To validate that mNmaA^M^ indeed contains an aspartimide moiety, we reacted 0.1 mM mNmaA^M^ with 2M hydrazine in 50 mM Tris-HCl buffer at pH 8. The reaction mixture was incubated at room temperature for 30 mins. If mNmaA^M^ contains an aspartimide, a mass shift of 32 Da is expected (Figure 4a,b). The incubation resulted in species with a +32 Da mass shift, confirming the presence of an aspartimide in mNmaA^M^.

**Figure 4:**
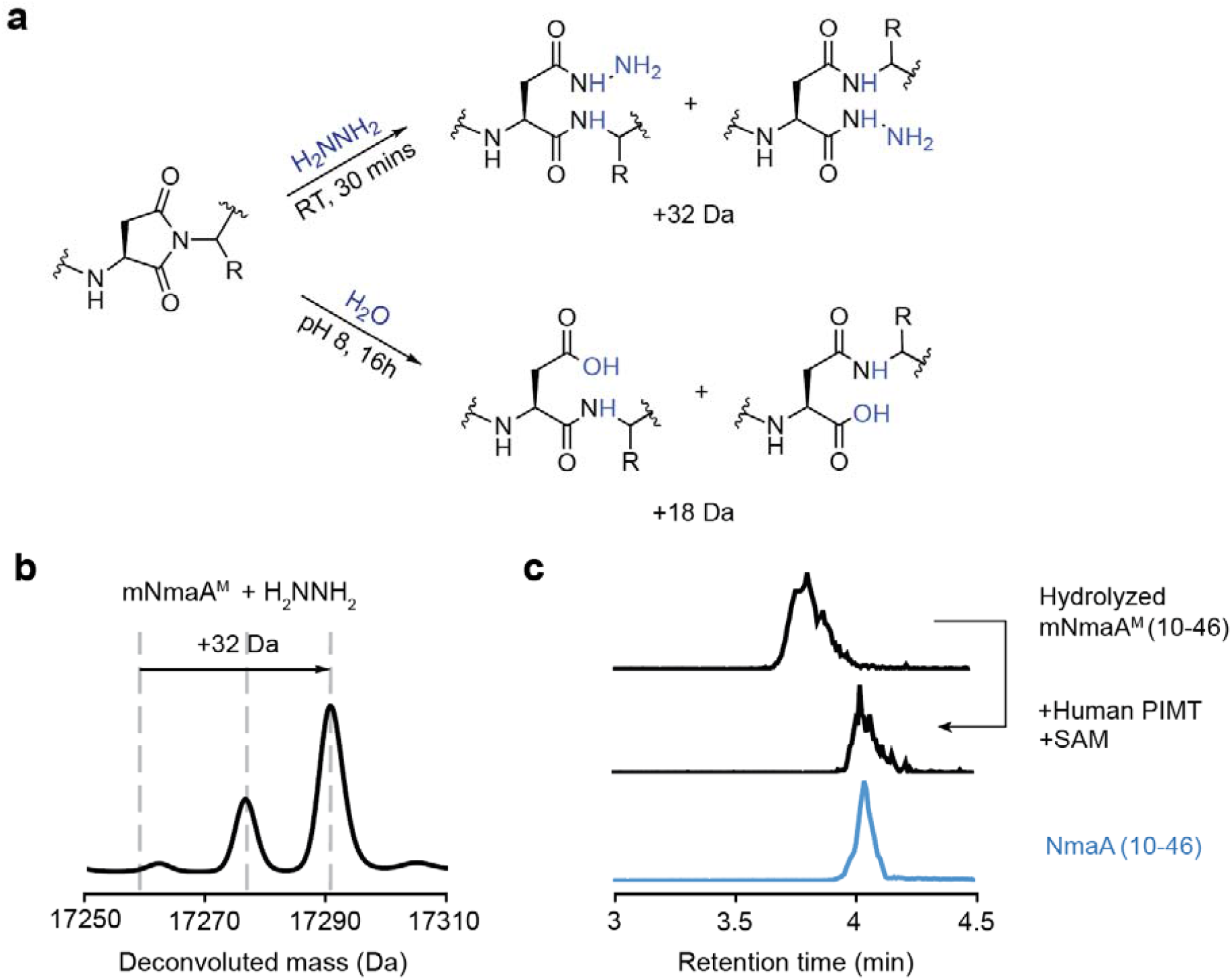
mNmaA^M^ Contains an Aspartimide Moiety. **a)** Scheme of hydrazinolysis and hydrolysis of aspartimide. Aspartimides can react with hydrazine at room temperature for 30 mins to form hydrazides, resulting a gain of 32 Da. In addition, aspartimides can also react with water in weak alkaline condition (pH 8), forming a mixture of aspartate and isoaspartate. **b)** Mass spectrum of mNmaA^M^ reacted with hydrazine. A +32 Da mass shift was observed upon reactions with hydrazine, demonstrating mNmaA^M^ contains aspartimide. **c)** Hydrolyzed mNmaA^M^ contains isoaspartate. EIC spectra of hydrolyzed mNmaA^M^ (10-46) and unmodified NmaA (10-46) are shown in black and blue, respectively. mNmaA^M^ (10-46) had a retention time of 3.8 min. After treating this sample with human PIMT and SAM, isoaspartate residue was reverted back to aspartate, as suggested by the same retention time as NmaA (10-46).

Aspartimides are known to be liable and can be hydrolyzed at neutral pH in buffer solutions (Figure 4a). Therefore, to investigate the hydrolyzed products of mNmaA^M^, we incubated it in 50 mM Tris-HCl, pH 8 at room temperature for 16h, and then digested the protein with endopeptidase GluC. This digestion generated a C-terminal 37-aa fragment with a missed GluC digest site at E31 in NmaA sequence (Figure 3c), which we refer to as hydrolyzed mNmaA^M^ (10-46). LC-MS analysis of this species revealed that the aspartimide was indeed hydrolyzed under this reaction condition. Moreover, the hydrolyzed mNmaA^M^(10-46) had a retention time of 3.8 min, while the GluC-digested unmodified His_6_-SUMO-NmaA control NmaA(10-46) had a retention time of 4 min (Figure 4c). This observation suggested that the aspartimide in mNmaA^M^ is converted regioselectively to form a β-amino acid, isoaspartate (Figure 4a). To confirm this hypothesis, we reacted this species with human PIMT *in vitro*, as human PIMT can revert isoaspartate residues back to aspartate. Indeed, the human PIMT recognized this species, fully converting the species to a retention time that matched with the NmaA(10-46) control, which only contained the proteinogenic α-amino acids (Figure 4c).

Next, we sought to determine the location of the aspartimide in mNmaA^M^. The NmaA sequence contains six Asp residues (Figure 3c). However, only 4 Asp residues can be aspartimidylated, as P8 and P18 lack the amide proton to form aspartimides with D7 and D17 sidechains, respectively. We first digested mNmaA^M^ with endoproteinase GluC, which generated a C-terminal 37-aa peptide bearing the modification, which we refer to as mNmaA^M^(10-46). Subjecting this peptide to MS/MS fragmentation (Figure S9) revealed that the modification was located between Gln22 and His29. However, because this region was located right in the middle of the fragment, the *b*/*y*-ion coverage was not sufficient for suggesting the exact location of the aspartimide. Therefore, to determine the exact location, we generated D23N, D25N, and D28N variants. Coexpressing these variants with NmaM suggested that neither D23N nor D25N variant affect the aspartimide formation (Figure S10). In comparison, substituting D28 to N resulted in no modification on the precursor, indicating that aspartimide formed between the sidechain of D28 and the backbone amide of H29.

To provide spectroscopic evidence of the aspartimide formation, we acquired the FTIR spectrum of the GluC digested mNmaA^M^ D23E variant. The mNmaA^M^ D23E variant can be modified as well as the wildtype substrate (Figure S11). The GluC digestion of the mNmaA^M^ D23E variant generates a 23 aa C-terminal fragment that can be reasonably purified using HPLC, which we denote this fragment as NmaA^M^ (24-46). The FTIR spectrum of NmaA^M^ (24-46) has an additional shoulder peak at 1708 cm^−1^ compared to the unmodified NmaA (24-46) control (Figure S12). To validate this peak corresponds to an aspartimide formation, we also acquired FTIR spectra of amycolimiditide and pre-amycolimiditide reported by our previous studies. Both amycolimiditide and pre-amycolimiditide have 4 ester linkages, but amycolimiditide has an additional aspartimide moiety suggested by the solution NMR structure (PDB: 8DYM).^21^ Indeed, both spectra showed a peak at 1742 cm^−1^, indicating the presence of esters. Additionally, the FTIR spectrum for amycolimiditide also contains an additional peak at 1708 cm^−1^, indicating this peak corresponds to an aspartimide formation (Figure S12).

### His29 is the Preferred Residue C-terminal to the Aspartimide in mNmaA^M^

Clarke and coworkers have collectively reported several experimental studies on aspartimide formation from aspartyl peptide, suggesting that the sidechain of the residue that contributes its backbone to form the aspartimide (i.e. the *n* + 1 residue) greatly affects the rate of aspartimide formation.^25,26^ In lasso peptides with an aspartimide modification, the *n* + 1 residue is highly conserved to be Thr, and our studies have shown that the *n* + 1 residue in these peptides plays a huge role in determining the rate of methylation and the subsequent aspartimide formation.^15^ Moreover, we have recently reported that the *n* + 1 residue, Thr7, in lasso peptide lihuanodin also controls the regioselectivity of the aspartimide hydrolysis and subsequently the threadedness of the lasso peptide.^16^ In comparison, the *n* + 1 residue for aspartimides in lanthipeptides and graspetides are predominantly Gly,^13,17,18^ which has been shown to be the preferred residue for aspartimide formation in both computational and experimental studies. ^21,27^

As illustrated above, in mNmaA^M^, the *n* + 1 residue is H29, which is uncommon in predicted imiditide precursors as well as other PIMT-associated RiPPs. To probe whether histidine can serve as an effective *n* + 1 residue for methylation and aspartimide formation in mNmaA^M^ biosynthesis compared to the well-characterized Thr and Gly, we carried out site mutagenesis at this position. We first constructed the H29T variant to mimic the aspartimidylation site seen in lasso peptides cellulonodin-2 and lihuanodin (Figure 5). Coexpressing this variant with NmaM at room temperature for 20h showed an increase in the accumulated methylated species compared to the wildtype peptide. This observation suggests that having Thr as the *n* + 1 residue in mNmaA^M^ leads to a fast methylation step, followed by a relatively slower aspartimide formation step. This observation is also supported by the rate measurements of aspartimide formation in lihuanodin.^16^ Nevertheless, the percentage of aspartimidylated product formed in H29T variant was lower than the wildtype under the same heterologous expression condition. Additionally, we constructed the H29G variant to resemble the aspartimides in lanthipeptides and graspetides. The coexpression of this variant with NmaM showed nearly no accumulation of the methylated species (Figure 5), suggesting that aspartimide formation happened readily as soon as the methylated species was formed. This result also agreed with the heterologous expressions of aspartimide-containing graspetides, where having Gly as the *n* + 1 residue led to no accumulations of methylated species.^17,21^ Similarly, the amount of aspartimidylated H29G variant was also lower compared to the wildtype peptide, suggesting that the H29G variant was not as good of a substrate for NmaM. These mutagenesis measurements collectively illustrated that H29 in wildtype mNmaA^M^ is the preferred *n* + 1 residue for its biosynthesis compared to the commonly adopted Thr or Gly. As the *n* + 1 residue for aspartimidylation has been constrained to mainly Gly and Thr in previous studies, this finding helps to expand the list of potential *n* + 1 residues for predicting the site of aspartimidylation bioinformatically.

**Figure 5:**
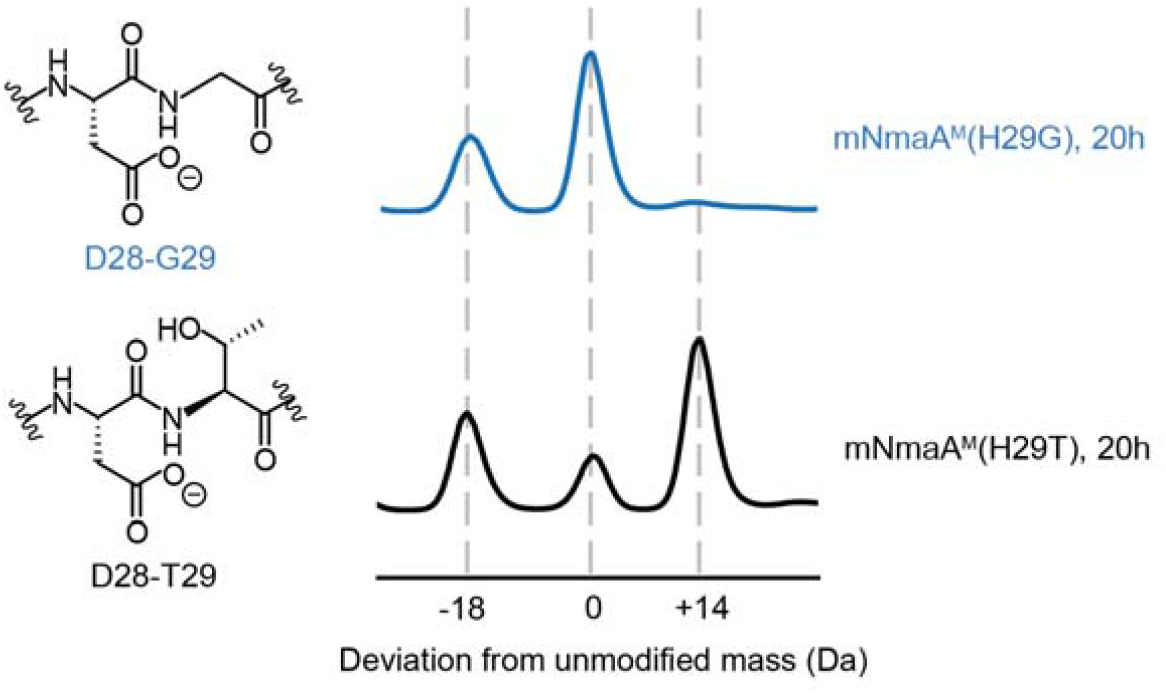
Effects of Single Amino Acid Substitutions of *n +* 1 Residue on Methylation and Aspartimidylation. Both mNmaA^M^ H29T and H29G variants were heterologously expressed under the same conditions as the wild-type substrate (20h). the H29T substation caused a larger buildup of the methylated species compared to the wild type, showing that the rate of aspartimidylation was slower than methylation. In addition, less aspartimidylated products were produced for the H29T variant compared to the wildtype. On the other hand, the H29G substitution led to no buildup in the methylated species, showing that rate of aspartimidylation was much faster than methylation when Gly was the *n* + 1 residue. Similarly, less aspartimidylated product was observed, showing the H29G variant was not as good of a substrate compared to the wildtype peptide. Collectively, His29 is the preferred *n* + 1 residue for aspartimidylation for NmaA.

### Much of Leader and Follower Peptides of mNmaA^M^ is Dispensable

One shocking aspect of the mNmaA^M^ heterologous expression is that the modified peptide binds strongly to the modifying enzyme NmaM even in the presence of 8M urea (Figure 3e). In addition, it is fascinating how NmaM achieves specificity for aspartimylating Asp28 on NmaA out of an abundance of aspartate residues *in vivo*. To address this question, we first attempted to decipher the elements on the precursor peptide that are important for substrate recognition. We first constructed variants with truncations of the NmaA leader peptides, removing 5 aa at a time, up to 20 aa in total (Figure 6a). The Met residue coded by the start codon was not included as the precursor was fused to a His_6_-SUMO tag. Coexpressing these variants with NmaM showed that removing the first 5 and 10 amino acid of the leader had no effect on aspartimidylation of NmaA (Figure 6b). Additionally, these variants could pull down NmaM as well as the full length NmaA, showing that the first 10 residues in the leader peptide were not important for substrate recognition (Figure S13). In contrast, a decrease in aspartimidylation of NmaA was noted when the first 15 aa of the leader peptide was removed. In the meantime, we also observed a decrease in amount of NmaM that the truncated variant could pull down (Figure S13), suggesting that this region was important for substrate recognition for NmaM. Removing a total of 20 aa of the leader peptide resulted in a more severe drop in aspartimidylated product formation and an even less amount of NmaM pulled down (Figure 6b, S13). It is worth noting that there is a high content of basic amino acids in the region of NmaA(12-21), which can play an important role in substrate recognition through electrostatic interactions. Nevertheless, the aspartimidylation could still occur even when the first 20 aa of leader peptide was removed, showing that much of the leader peptide of mNmaA^M^ is dispensable.

**Figure 6:**
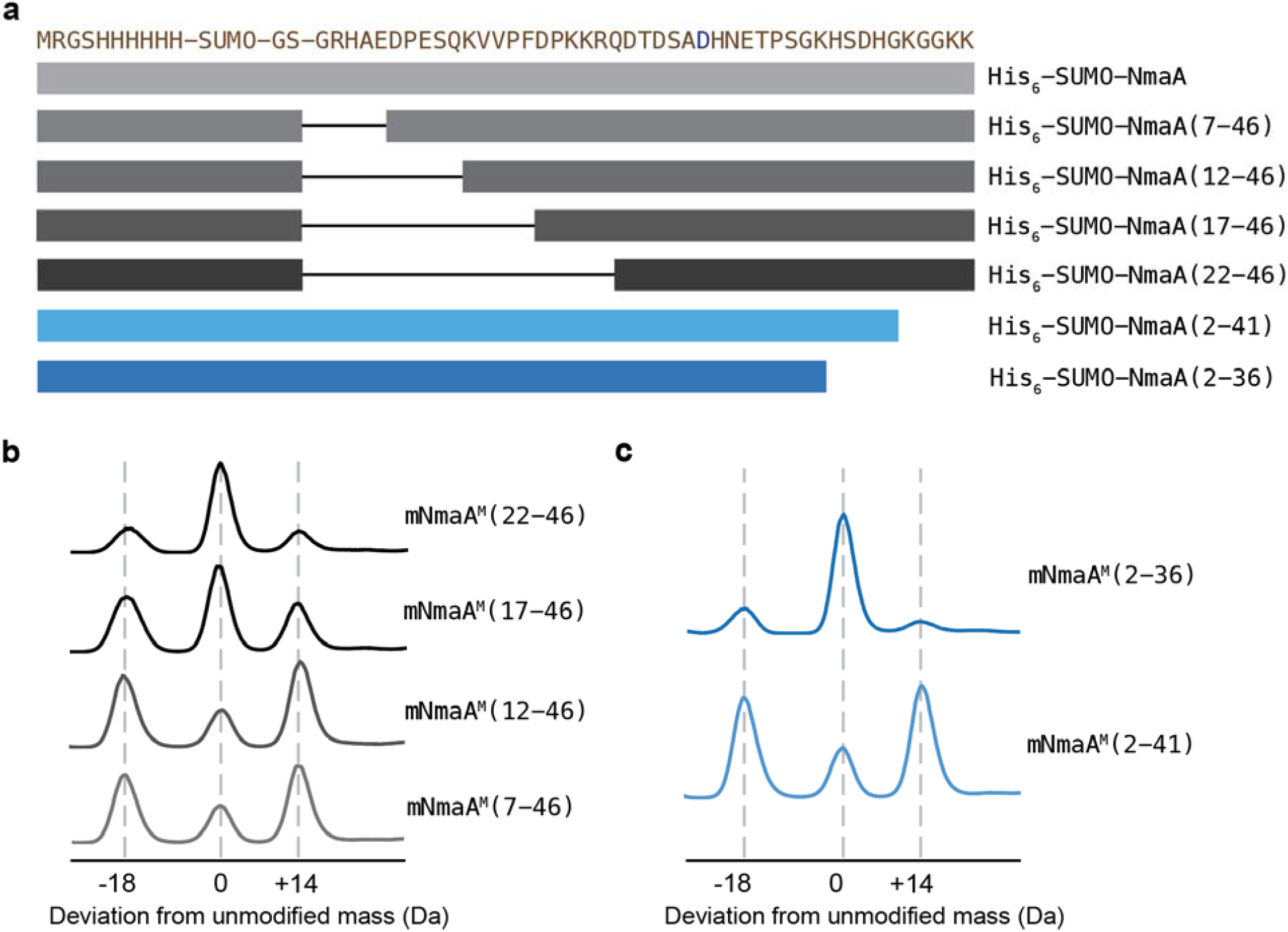
Effects of Leader and Follower Truncations on mNmaA^M^ Biosynthesis. **a)** Sequence of His_6_-SUMO-NmaA and schematic of the leader and follower truncation variants. The site of modification, Asp28, is colored blue. **b)** Much of the leader is dispensible for mNmaA^M^ biosynthesis. 5 aas are removed at a time. Removing the first 10 aa in the leader had minimal effect on aspartimidylation. Removing the next 10 aa was detrimental to the substrate recognition (Figure S13) and the extent of aspartimidylation, likely due to the removal of basic amino acids in this region. **c)** Much of the follower is also dispensible for mNmaA^M^ biosynthesis. Removing the first 5 aa on the follower sequence had minimal effect on aspartimidylation. Removing the next 5 aa significantly reduced the extent of the reaction, showing this region was important for substrate recognition.

In some cases of RiPP biosynthesis, such as bottromycin^28,29^ and cyanobactin,^30,31^ a C-terminal follower sequence to the core peptide has been shown to be important for RiPP maturation. Since the modification on mNmaA^M^ occurs in the middle of the precursor sequence, we set to ask whether a C-terminal recognition sequence exists for imiditide biosynthesis. In a similar fashion, we constructed variants with 5 aa and 10 aa truncations of the precursor from the C-terminus (Figure 6a). Removing the first 5 aa from the C-terminal domain had minimal effect on aspartimidylation of NmaA (Figure 6c). However, we observed a decrease in the amount of NmaM pulldown compared to the full length NmaA (Figure S14), suggesting that the basic lysine residues near the C-terminus could also be important for binding between NmaA and NmaM. In comparison, removing the next additional 5 aa from the C-terminus was detrimental to the extent of aspartimidylation on NmaA (Figure 6c), but had no effect on the amount of NmaM getting pulled down (Figure S14). Therefore, our data suggest that much of the follower sequence is also dispensable for aspartimidylation to occur in the mNmaA^M^ biosynthesis. Nevertheless, both the leader and follower sequences were crucial for the strong binding between NmaA and NmaM, demonstrating how the reaction specificity of NmaM was achieved *in vivo*.

### Importance of the C-terminal Domain of NmaM in Substrate Recognition

Our methodology of genome mining suggests that the *O*-methyltransferases that carry out class-defining aspartimidylation in imiditide BGCs share conserved C-terminal domains that are absent in canonical PIMTs (Figure 2a, S2). In our previous work on the PIMT homologues that aspartimidylate lasso peptides, we showed that this C-terminal extension to the canonical PIMT is important for the substrate recognition.^15^ Therefore, to examine the importance of the identified C-terminal motif (Figure 2a) in substrate recognition, we first constructed the 50 aa C-terminal truncated variant of NmaM (NmaMΔC50) that lacked this motif. We coexpressed NmaMΔC50 with His_6_-SUMO-NmaA, and observed no modification on the precursor (Figure S15). Meanwhile, His_6_-SUMO-NmaA was no longer able to pull down this variant (Figure S15), indicating that the C-terminal motif was essential for substrate recognition.

In addition to the conversed motif, we noticed that the ∼150 aa C-terminal domain of NmaM is rich in negatively charged residues and has an overall pI value of 5.4 (Figure S16). As demonstrated above, the leader and follower peptides in the precursor peptide are rich in positively charged residues, and removing these residues is detrimental to substrate recognition (Figure 6b). Therefore, we hypothesized that the substrate recognition is likely achieved through charge-charge interactions. To test this hypothesis, we first identified a region in C-terminal domain of NmaM that has a very concentrated presence of negatively charged residues, namely DEDGD from position 289-293 in NmaM. Then, we constructed a NmaM variant where this region was substituted with SGSGS to remove 4 negative charges from the C-terminal domain, which we named NmaM(SGSGS). Coexpressing the NmaM(SGSGS) variant with His_6_-SUMO-NmaA resulted in predominately unmodified His_6_-SUMO-NmaA, indicating that these negative charged residues in NmaM are important for mNmaA^M^ maturation (Figure S17). Additionally, the SDS-PAGE gel of the purified His_6_-SUMO-NmaA fractions suggested that mNmaA^M^ pulled down less NmaM(SGSGS) compared to the wildtype NmaM, supporting our hypothesis that the charge-charge interactions between NmaA and NmaM are crucial for substrate recognition (Figure S17).

### AlphaFold Model Predicts Mode of Substrate Recognition by NmaM

To further understand the nature of the substrate specificity of NmaM on NmaA, we generated an AlphaFold model of the NmaA-NmaM complex using ColabFold,^32,33^ assuming a 1:1 stoichiometry between the two proteins (Figure 7a, b). To our surprise, the model near perfectly supports our experimental results. Even though the precursor NmaA is predicted to be a random coil, various hydrogen bonding as well as charge-charge interactions ensure the strong binding and substrate specificity between NmaA and NmaM (Figure S18). The side chain of Asp28 sits right outside of the predicted SAM binding pocket in NmaM, suggesting that Asp28 is the site of modification (Figure 7c). The C-terminal domain of NmaM binds to both the leader and core portion of the leader peptide, supporting this additional C-terminal domain compared to canonical PIMTs in NmaM is essential for substrate recognition (Figure 7b). Additionally, the N-terminal 50 aa domain that is absent in canonical PIMTs seems to be involved in binding to the core peptide (Figure 7b). The side chain of the *n* + 1 residue His29 appears to interact with F284 in NmaM through CH−π interaction (Figure S19),^34^ further promoting the substrate specificity. For NmaA leader peptide, the model suggests that NmaA(1-10) doesn’t not interact with NmaM, agreeing with our experimental result that removing NmaA (1-10) had minimal effect on the mNmaA^M^ maturation. On the other hand, NmaA(11-20) binds to the C-terminal domain of NmaM, and experimental results showed removing NmaA (11-20) had deleterious effect on the extent of modification on NmaA. In contrast, the model indicates the follower sequence doesn’t interact with NmaM. However, based on experimental results above, we hypothesized that the follower sequence that is rich in negatively charged residues should be engaged with the acidic patch DEDGD at NmaM (289-293), as they are spatially adjacent based on the model (Figure S20).

**Figure 7:**
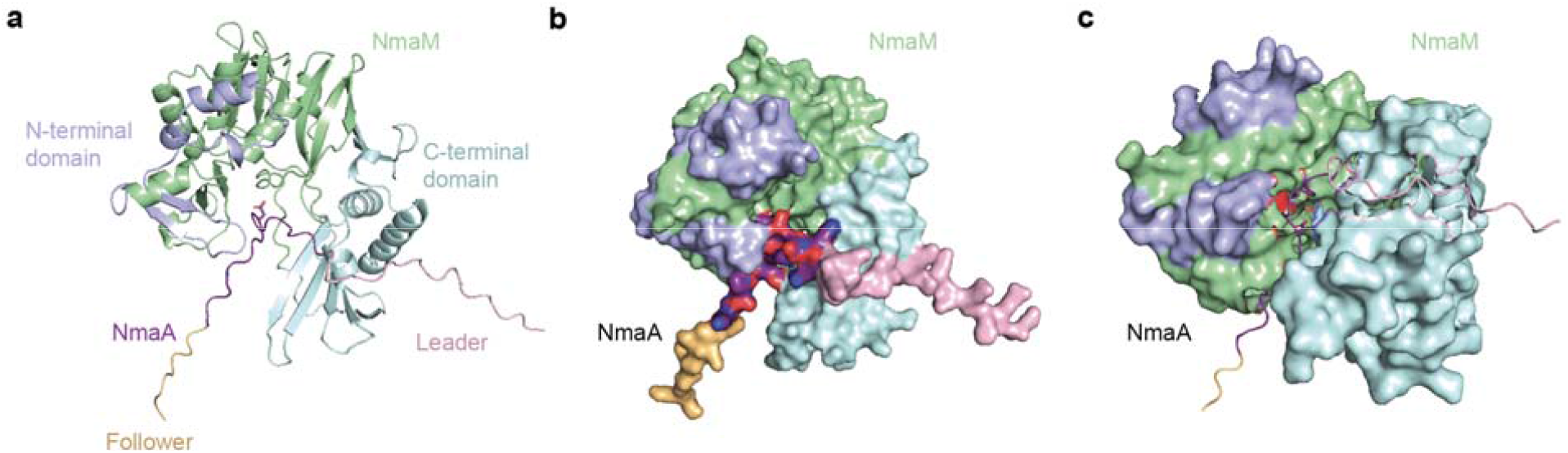
AlphaFold Model of NmaA-NmaM using ColabFold. ^32,33^. **a)** AlphaFold model of NmaA-NmaM complex. A stoichiometry of 1:1 between NmaA and NmaM is assumed. The leader, core, and follower peptide of NmaA are colored pink, purple and gold, respectively. The N-terminal domain and the C-terminal domain of NmaM are colored lightblue and cyan, respectively. The rest of NmaM is colored green. The sidechain of Asp28 is shown in stick. **b)** Space filling model of Nma-NmaM. The leader peptide binds to the C-terminal domain of NmaM. The core sequence is locked in near the active site of NmaM. The follower sequence was not engaged with NmaM based on the model, but our experimental data suggest that the follower peptides also bind with the C-terminal domain of NmaM. **c)** Asp28 sits right outside of the SAM binding pocket of NmaM (red), suggesting that it is the site of modification.

Considering how well the AlphaFold prediction model agrees with our experimental data acquired for the mNmaA^M^ biosynthesis, we advocate that AlphaFold can serve as a powerful tool guiding novel RiPPs discovery. The putative precursor peptide can be readily tested to see whether they can serve a substrate for nearby encoded putative RiPP enzymes to high accuracy when compared to experimental results based on previous studies.^35^ Additionally, AlphaFold can also help indicate which reaction happens first when several enzymes are involved in the biosynthetic gene cluster (Figure S21).^17,35^

## Discussion

In this work we have described the discovery and the genome mining of a novel family of RiPPs, the imiditides. Utilizing the fact that PIMT homologs install the aspartimide moiety on peptides from various RiPP families, we used the conserved motif on the extended C-terminal domain of these PIMT homologs as bioinformatic seeds to find this novel family of RiPPs. Our bioinformatic analysis suggested that imiditides are widely distributed in Gram-positive bacterial genomes. In addition, we reported the discovery of the founding member of the imiditide family, mNmaA^M^, from *Nonomuraea maritima*. The aspartimide in mNmaA^M^, is unusually stable, which accumulates and is the major product in the heterologous expression in *E. coli*. In contrast to other RiPP-associated PIMT homologs that recognize constrained peptides as substrates (i.e. lasso peptides, grasepetides, lanthipeptides),^13,17,21,36^ imiditide-associated PIMT homologs recognize specific Asp residues on the linear precursor. In mNmaA^M^ biosynthesis, the modifying enzyme NmaM methylates a specific Asp residue in the precursor sequence NmaA through extensive charge-charge interactions and hydrogen bonding networks. This specificity is crucial for the fitness of the host as random methylation on Asp residues will affect functions of numerous proteins and constantly consume the precious SAM molecules.

The discovery of imiditides challenges the current understanding of how aspartimide can be introduced in the cells.^37^ Aspartimides have been long considered as the unstable intermediates resulted from spontaneous protein aging and degradation.^38–40^ However, our finding suggests the PIMT homologs in imiditide BGCs install the aspartimide intentionally on linear peptides. Therefore, future discovery of aspartimide moieties in the Gram-positive bacterial proteomes should also consider the existence of imiditides rather than solely regarding them as protein degradation byproducts. The persistence of aspartimide in imiditides depends on their final disposition of peptides. In extracellular environments, in the case of mNmaA^M^, the aspartimide can be hydrolyzed regioselectively to isoaspartate. On the other hand, if the aspartimide persists, it can serve as an electrophile for further modifications.

Moreover, we are fascinated by the AlphaFold prediction of NmaA-NmaM complex, which nearly perfectly predicts our experimental results. It not only predicts that NmaA will be a substrate to NmaM, but also correctly predicts the site of modification. Other than the fact that the follower sequence should interact with the acidic loop in the C-terminal domain of NmaM based on our experimental results (Figure S20), it predicts the binding interface between NmaA and NmaM extremely well. We used AlphaFold as a validation tool in this study since much of our experimental results were gathered before the release of ColabFold. However, we argue that AlphaFold can be a powerful tool in predicting whether a biosynthetic gene cluster can form a RiPP, and help design experiments in validating the gene cluster. In summary, we have discovered a novel RiPP family where the aspartimide formation is the class-defining modification. Future studies will be directed towards the discovery of other novel RiPPs with the aspartimide modification, as well as understanding the role of aspartimides in the bioactivity of these RiPPs.

## Acknowledgments

This work was supported by the National Institutes of Health Grant GM107036 and a grant from Princeton University School of Engineering and Applied Sciences (Focused Research Team on Precision Antibiotics). L.C. was supported by an NSF Graduate Research Fellowship Program under Grant DGE-1656466 and a Proctor Fellowship from Princeton University.

## Notes

### Competing Interest Statement

The authors have declared no competing interest.

