## Supporting Info for "Genome Mining and Discovery of Imiditides, a Novel Family of RiPPs with a Class-defining Aspartimide Modification"

### Supplementary Materials and Methods

**Materials and Equipment**

*Escherichia coli* XL-1 Blue were used for all recombinant DNA procedures. All primers were ordered from Integrated DNA Technologies (IDT). *nmaAM* gene cluster was codon optimized for *Escherichia coli and* assembled using primer annealing and gBlocks HiFi Gene Fragments from IDT. Restriction enzymes, Q5 DNA polymerase, and T4 DNA ligase were purchased from New England Biolabs (NEB). The sequences of all primers used are provided in Table S1. All sequencing results were confirmed by Sanger sequencing (Genewiz/Azenta). Acetonitrile was purchased from Sigma-Aldrich. Semi-preparative reverse-phase HPLC was performed using a Zorbax 300SB-C18 (9.4 mm x 250 mm, 5 μm) column with an Agilent 1200 series instrument. LC-MS and LC-MS/MS experiments were done using a Zorbax 300SB-C18 (2.1 mm x 50 mm, 3.5 μm) column with a 1260 Infinity II system coupled to an Agilent 6530 qTOF instrument. Infrared spectra were collected on a Bruker VERTEX 80v spectrometer equipped with a liquid nitrogen cooled mercury cadmium telluride (MCT) detector.

**Cloning**

The gene containing *nmaA* was codon optimized for *Escherichia coli* XL-1 and assembled using primer annealing, digested using *Nco*I and *Hin*dIII restriction enzymes, and ligated into first multiple cloning site of pRSF-Duet. The *nmaA* was further amplified and fused to a SUMO-tag, digested using *Bam*HI and *Hin*dIII restriction enzymes, and ligated into pQE-80, creating pLC53. Genes encoding SUMO-NmaA variants were cloned using pLC53 as a template, and mutations or truncations were introduced on primers. The resulting fragments were digested using *Bam*HI and *Hin*dIII restriction enzymes, and ligated to pQE-80.

The gene containing *nmaM* was also codon-optimized for *E. coli*, and was amplified from IDT gBlocks HiFi Gene Fragments. It was then digested using *Nde*I and *Avr*II restriction enzymes, and ligated into the second multiple cloning site of pRSF-Duet, forming pLC55. For the construct that expressed NmaM constitutively, *nmaM* was placed under a constitutive *mcjBCD* promoter from the microcin J25 gene cluster was appended upstream of *nmaM* in an overlap PCR. This PCR product was then cloned in the forward direction into pLC53 using *Nhe*I and *Nco*I restriction sites. Genes encoding NmaM variants were cloned using pLC55 as a template, and mutations or truncations were introduced on primers. The resulting fragments were digested using *Nde*I and *Avr*II restriction enzymes, and ligated to pRSF Duet.

**Expression and Purification of His_6_-SUMO-NmaA and Its variants**

The plasmid containing the N-terminal 6xHis tagged SUMO-NmaA was transformed into *E. coli* BL21(DE3) *ΔslyD* cells or *E. coli* BL21(DE3) *ΔslyD Δpcm* and grown up overnight in LB with 100 mg/L ampicillin. For coexpression with NmaM, the plasmid containing the untagged NmaM was also transformed. The overnight culture in LB will also contain 50 mg/L kanamycin. The cells were subcultured at an OD_600_ = 0.02 into 500 mL of LB with 100 mg/L ampicillin (or also with 50 mg/L kanamycin for coexpression with NmaM) and allowed to grow at 37 °C. Once the OD_600_ = 0.5, expression was induced using 1 mM IPTG, also at 37 °C. The cells were grown for 20 hours at 20 °C before being spun down at 4000 x *g*. Proteins are purified either using the denaturing condition or the native condition.

For denaturing purification, the pellet from a 500 mL culture was resuspended in 10 mL 100 mM NaH_2_PO_4_, 10 mM Tris, 8 M urea, pH 8.0. After freezing at -80 °C for 30 mins and thawing at room temperature, the cell lysate was centrifuged at 8000 x g for at least 20 minutes. The clarified lysate was mixed with 1 mL Ni-NTA resin (Qiagen) and incubated while rotating at 4 °C for 1 hour. The mixture was passed through an empty gravity column, followed by the first wash with 10 mL 100 mM NaH_2_PO_4_, 10 mM Tris, 8 M urea, pH 6.3, and the second wash with 10 mL 100 mM NaH_2_PO_4_, 10 mM Tris, 8 M urea, pH 5.9. Then, the protein was eluted with a total of 8 mL of 100 mM NaH_2_PO_4_, 10 mM Tris, 8 M urea, pH 4.5, collecting 1 mL fraction at a time. After analyzing the fractions with SDS-PAGE, the clean elution fractions were pooled, buffer exchanged into 50 mM Tris-HCl, 100 mM NaCl, 10% glycerol, pH 8, and concentrated using a 10 kDa Amicon concentrator. The purified proteins were stored at -80 °C.

For native purification, the pellet from a 500 mL culture was resuspended in 10 mL of 50 mM NaH_2_PO_4_, 300 mM NaCl, 10 mM imidazole pH 8.0. The sample was then sonicated on ice to lyse the cells and spun down at 4000 x *g* for 10 minutes at 4 °C. The supernatant was removed and spun down an additional time at 8000 x *g* for 10 minutes. The clarified lysate was then incubated with 1 mL of Ni-NTA resin (Qiagen) while rotating for 1 hour. The mixture was added to an empty gravity column, and the flowthrough was passed over the resin an additional time. The resin was then washed with 10 mL of 50 mM NaH_2_PO_4_, 300 mM NaCl, 20 mM imidazole pH 8.0 and twice with 10 mL of 50 mM NaH_2_PO_4_, 300 mM NaCl, 50 mM imidazole pH 8.0. The protein was then eluted with 50 mM NaH_2_PO_4_, 300 mM NaCl, 250 mM imidazole pH 8.0. The samples were run on SDS-PAGE and the most concentrated samples were pooled and buffer exchanged using a PD-10 desalting column into 50 mM Tris-HCl, 100 mM Tris, pH 8.0. Purified and buffer exchanged protein were frozen in 10% glycerol and stored in aliquots and stored at -80 °C.

**GluC Digestion of His_6_-SUMO-NmaA and Its Variants**

GluC digestion of His_6_-SUMO-NmaA and its variants composed of 1 mg of protein substrate, 500 ng of GluC in 50 mM Tris-HCl, 2M urea, pH 8, for a total final volume of 200 μL. The reaction was carried out at 37 °C for 3 hours, before getting quenched using 20 μL of 10% formic acid. The reaction mixture was subsequently analyzed using LC-MS.

**HPLC of the GluC-Digested Fragments**

Typically for RP-HPLC purification, 25 μL of GluC digested SUMO-NmaA and its variant was injected onto the semi-preparative column. The mobile phase A consisted of water with 0.1% trifluoroacetic acid and phase B consisted of acetonitrile with 0.1% trifluoroacetic acid. The gradient program used was as follows: 5 % B from 0-1 minute, linear gradient from 10-45% B from 1-20 minutes, linear gradient from 45-90% B from 20-25 minutes, and isocratic elution at 90% B from 25-27 minutes.

**LC-MS and LC-MS/MS analysis of His_6_-SUMO-NmaA and Its Variants**

5 μL of purified His_6_-SUMO-NmaA and its variants were injected onto the XBridge Protein BEH C4 column (2.1mm x 50 mm, 300 Å, for proteins) or Zorbax 300SB-C18 (2.1 mm x 50 mm, 3.5 μm, for GluC digested fragments) column or with an Agilent 1260 Infinity II system for separation. The mobile phase A consisted of water with 0.1% formic acid, and phase B consisted of acetonitrile with 0.1% formic acid. The gradient used was: 10% B from 0-1 minute, linear gradient from 10-50% B from 1-20 minutes, linear gradient from 50-90% B from 20-25 minutes, and isocratic elution at 90% B from 25-30 minutes for proteins, and 5% B from 0-1 minute, linear gradient from 5-45% B from 1-20 minutes, linear gradient from 45-90% B from 20-25 minutes, and isocratic elution at 90% B from 25-30 minutes for peptides. The separated species were directly sprayed to an Agilent 6530 q-TOF instrument for mass detection and fragmentation.

**Hydrazine Reaction**

In the reaction mixture, modified SUMO-NmaA were added to 50 mM Tris-HCl to a final concentration of 0.1 mM and hydrazine (from a solution of 35% hydrazine in water, Sigma-Aldrich) was added to a final concentration of 2 M, with a final volume of 50 μL. The pH of the reaction mixture was set to 8. The reaction mixture was incubated at room temperature for 30 mins before getting analyzed by LC-MS.

***E. coli* BL21(DE3) *ΔslyDΔpcm* Construction**

*E. coli* BL21(DE3) *ΔslyD::kan* was first generated by P1 transduction from the Keio collection, where the donor strain was E. coli K12 MG1655 *ΔslyD::kan.* *kan* signified kanamycin-resistance marker. The recipient strain *E. coli* BL21(DE3) *Δpcm::kan* was confirmed by colony PCR. Then pCP20 plasmid was used to flip out the *kan* resistance marker, resulting *E. coli* BL21(DE3) *ΔslyD*. *E. coli* BL21(DE3) *ΔslyD Δpcm::kan* was further generated by P1 transduction from the Keio collection, where the donor strain was E. coli K12 MG1655 *Δpcm::kan.* The recipient strain *E. coli* BL21(DE3) *ΔslyD* *Δpcm::kan* was confirmed by colony PCR. pCP20 plasmid was further used to flip out the *kan* resistance marker, resulting *E. coli* BL21(DE3) *ΔslyD Δpcm*.

**Attenuated total reflectance Fourier transform infrared (ATR-FTIR) spectroscopy**

To prepare samples for ATR-FTIR analysis, lyophilized peptides were redissolved in ~ 500 μL ultrapure water (obtained from a Milli-Q system, EMD Millipore). Aliquots of 1 μL solution were added onto the surface of a diamond ATR crystal, which were dried under moisture- and CO_2_-free air. This step was repeated 10–15 times until a thin film of peptides was formed. The sample spectrum was measured as the average of 2,000 scans at a scan velocity of 80 kHz with a 5 mm aperture setting. A blank spectrum was collected immediately before each sample spectrum and automatically subtracted. Baseline corrections were performed on the ATR spectra after data collection.

**Genome Mining of Imiditides**

A set of 55 PIMT homologs identified by sequence similarity network analysis to be encoded within a minimal BGC having a RiPP precursor peptide sequence as the only other gene in the BGC were aligned with Multiple EM for Motif Elicitation (MEME), yielding a 41 aa sequence motif. The consensus sequence then served as the query sequence for protein BLAST analysis against the NCBI non-redundant protein BLAST database (https://ftp.ncbi.nlm.nih.gov/blast/documents/blastdb.html). Protein BLAST search was carried out using a standalone BLAST+ installation (package version 2.10.0) on the Princeton Della server.^1^ Using a relaxed e-value threshold of 1000, 5839 hits were obtained as potential PIMT homologs containing the sequence motif. The Genbank file in .gbff format for each organism predicted to encode a PIMT homolog was downloaded from NCBI and then uploaded to the Della server search for RiPP open-reading frames proximal to the PIMT coding sequence.

The next step of analysis was carried out using the custom Python script RiPP-scanner.py that can be found at the Link lab Github page. For each PIMT hit, the corresponding GenBank file for the host organism was loaded and the sequence information for the PIMT was retrieved from within the file using its RefSeq ID. The script then extracted the genomic sequence encompassing the PIMT sequence using user-defined size windows. This genomic region was then translated in both the forward and reverse directions, with 3 reading frames for each direction, using the NCBI translation table 11 for bacterial, archaeal, and plant plastid codons (https://www.ncbi.nlm.nih.gov/Taxonomy/Utils/wprintgc.cgi#SG27). Short open-reading frames (ORFs) with an initiating methionine or alternative start residues were identified from the translated sequences. As multiple short ORFs might be found near a PIMT, the final step in the analysis was to filter out false-positive hits that were unlikely to be true imiditide-encoding RiPP ORFs based on experimental observations of PIMT-associated imiditide RiPPs. The RiPP ORFs were required to be found within 500 base pair upstream or 1000 bp downstream of the PIMT homolog, facing the same direction as the PIMT gene. The putative imiditide was required to be 30 to 70 aa in length and contain at least one Asp residue for the modification. If multiple short ORFs met these filtering criteria, the ORF encoding a sequence most similar to NmaA, an experimentally verified imiditide, was identified and the remaining ORFs discarded based on pairwise alignment using the Biopython Bio.pairwise2 module. After applying these filters, 1002 imiditide hits were identified. To further reduce false positive hits, only imiditide hits associated with PIMT homologs that are 300-450 aa in length and that contain a GXGXG motif signifying a *S*-adenosyl methionine (SAM) binding site were retained. The GXGXG motif was also required to be found upstream of the 41 aa sequence motif identified by MEME. After applying these additional filters, 670 imiditide hits were identified.

**Sequence Similarity Network**

The sequence similarity network was generated using the enzyme similarity tool (EFI).^2^ ThfM protein sequence was used as the query sequence, and the maximum number of sequences retrived was set to 2000. UniProt sequence database was used for BLAST. An alignment score of 95 was used for generating the SSN. The network was visualized and analyzed using Cytoscape.

**Multiple Em for Motif Elicitation (MEME)**

The conserved C-terminal motif was generated using Multiple Em for Motif Elicitation (MEME) version 5.5.0.^3^ The motif discovery mode was set to classic mode. The number of motifs that MEME tried to find was set to 3. The list of proteins used for generating the motifs is listed in Table S1.

### Supplementary Figures

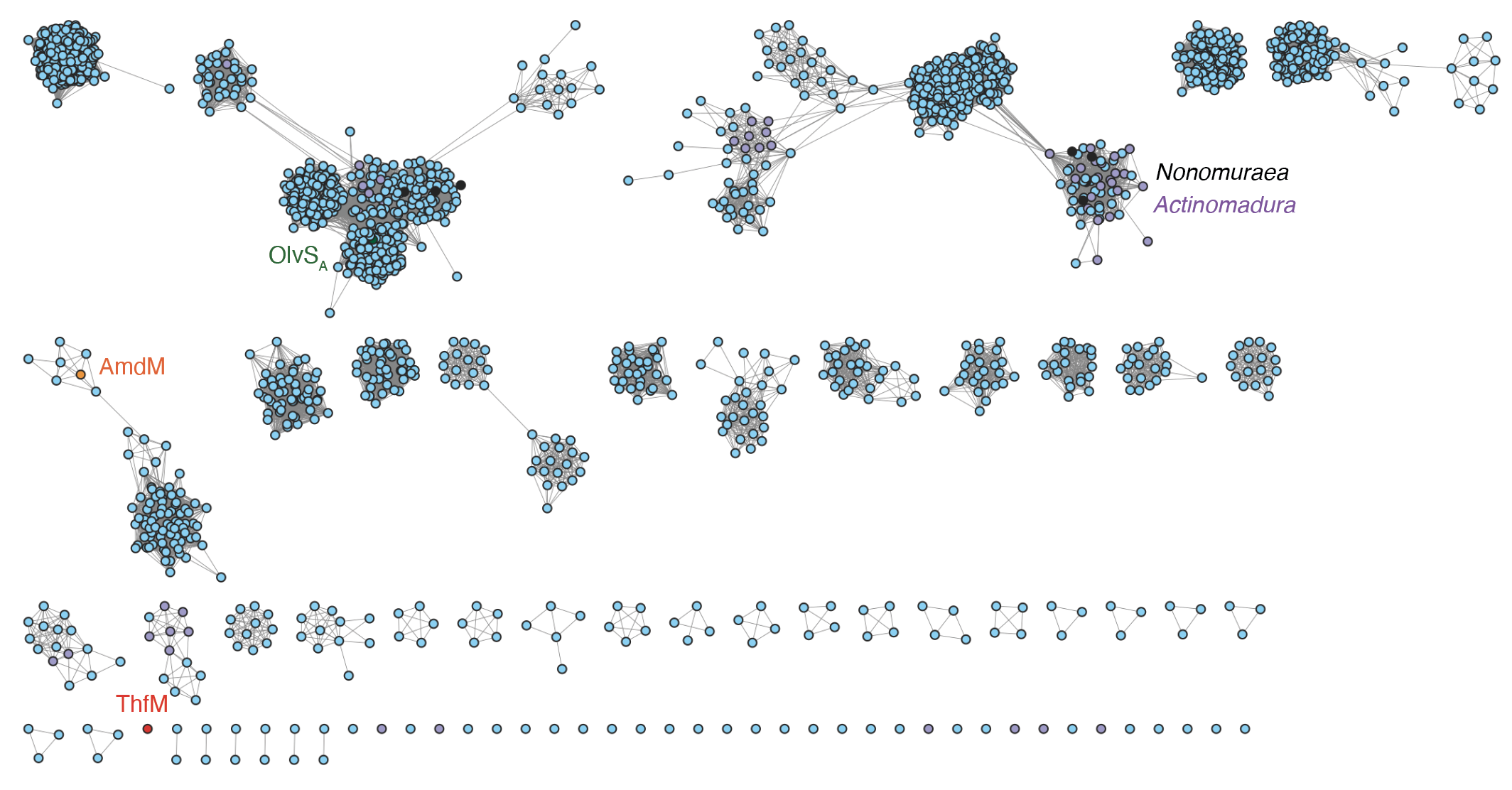

**Figure S1: Sequence Similarity Network (SSN) of ThfM.** 2000 sequences were queried in total, and an alignment score of 95 was used to create the sequence similarity network. ThfM does not cluster with any other sequence. AmdM clusters with several other methyltransferases from phylogenetically relevant strains that are also near graspetide biosynthetic gene clusters (BGCs). OlvS_A_ is in one of the major clusters with other methyltransferases near lanthipeptide BGCs. We identified a cluster that are rich in methyltransferases from *Nonomuraea* and *Actinomadura* strains. There are short open reading frames (ORFs) that cluster with these methyltransferases, which we hypothesize that these genes can constitute BGCs for a new family of RiPPs, imiditides.

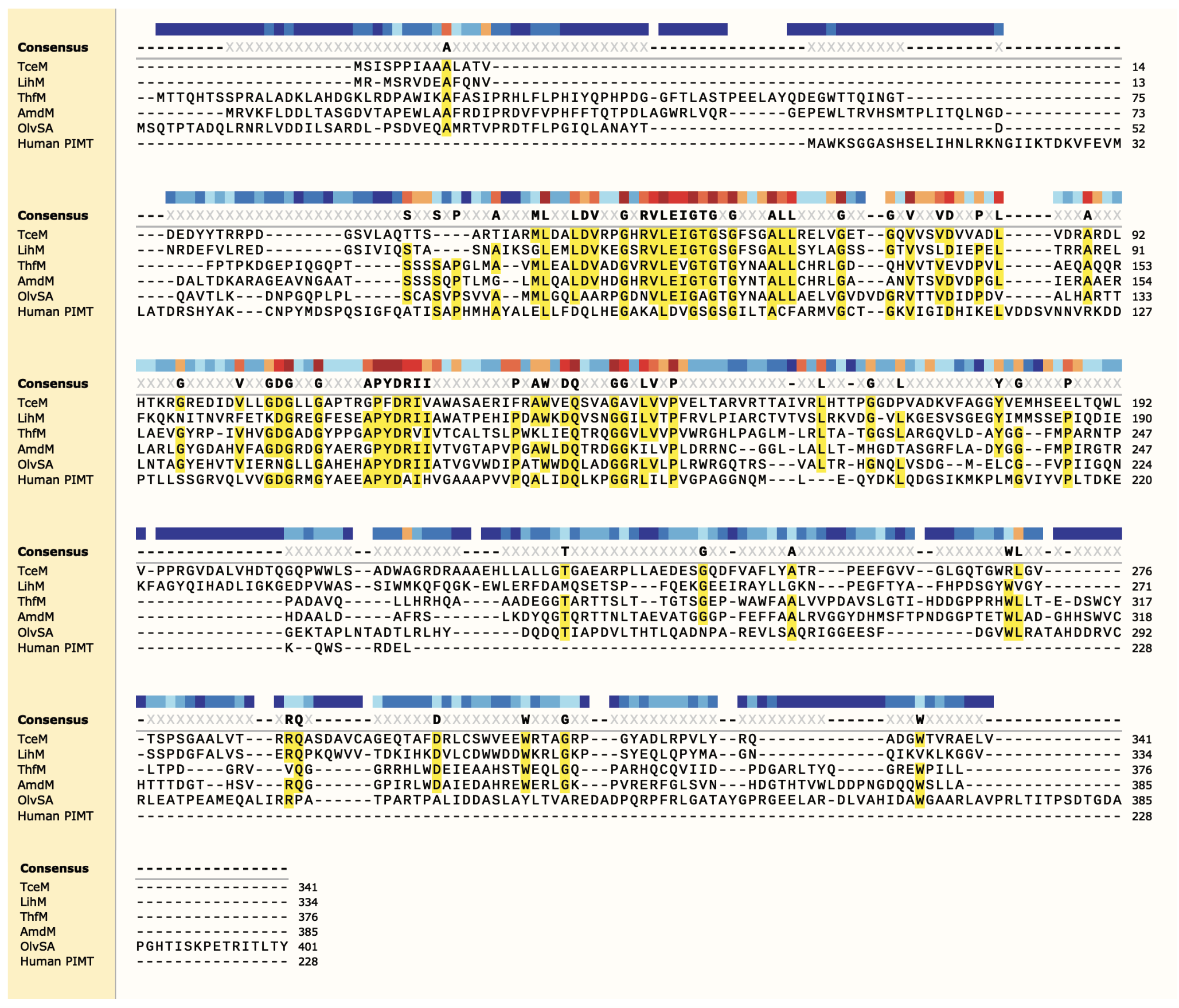

**Figure S2: Sequence Alignment of Methyltransferases Associated with Different Families of RiPPs.** TceM and LihM are PIMT homologs associated with lasso peptide biosynthetic gene clusters encoding cellulonodin-2 and lihuanodin.^4^ ThfM and AmdM are PIMT homologs associated with graspetide biosynthetic gene clusters encoding fuscimiditide and amycolimiditide.^5,6^ OlvS_A_ is a PIMT homolog associated with lanthipeptide OlvA(BCS_A_) biosynthetic gene cluster.^7^ Graspetide and lanthipeptide associated PIMT homologs have a ~50 aa N-terminal domain that are absent in canonical PIMT. All RiPP associated PIMTs have an additional C-terminal domain compared to canonical PIMT. However, the C-terminal domain sequences show more similarities when they are associated with the same family of RiPPs. Supported by ThfM SSN (Figure S1), the sequences of the C-terminal domain parts in graspetide and lanthipeptide associated methyltransferases have more resemblance compared to that in PIMT homologs associated with lasso peptides.

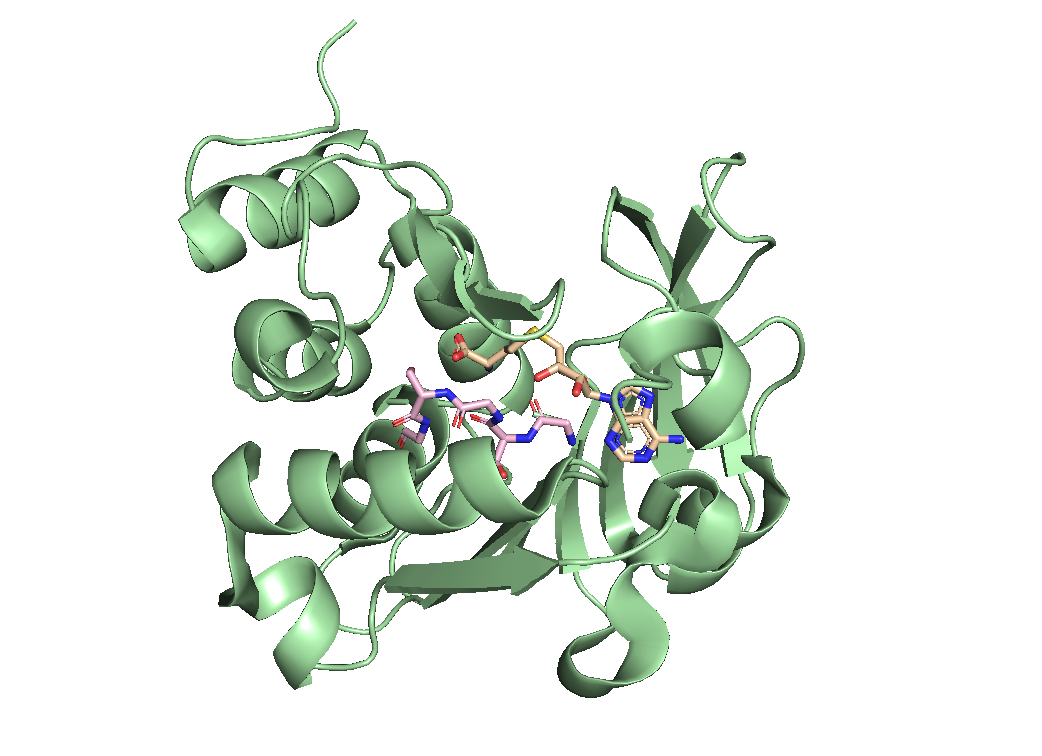

**Figure S3: Crystal Structure of Human PIMT with *S*-adenosyl Homocysteine (SAH)** (PDB: 1I1N). SAM was the methyl donor for the reaction. The reaction product SAH was used in solving the crystal structure and colored in wheat. The conserved GXGXG motif in PIMT family proteins (GSGSG in sequence of human PIMT) was colored in pink and shown in stick. The structure showed that the GXGXG motif bound to the amino part of SAM when the reaction was catalyzed.

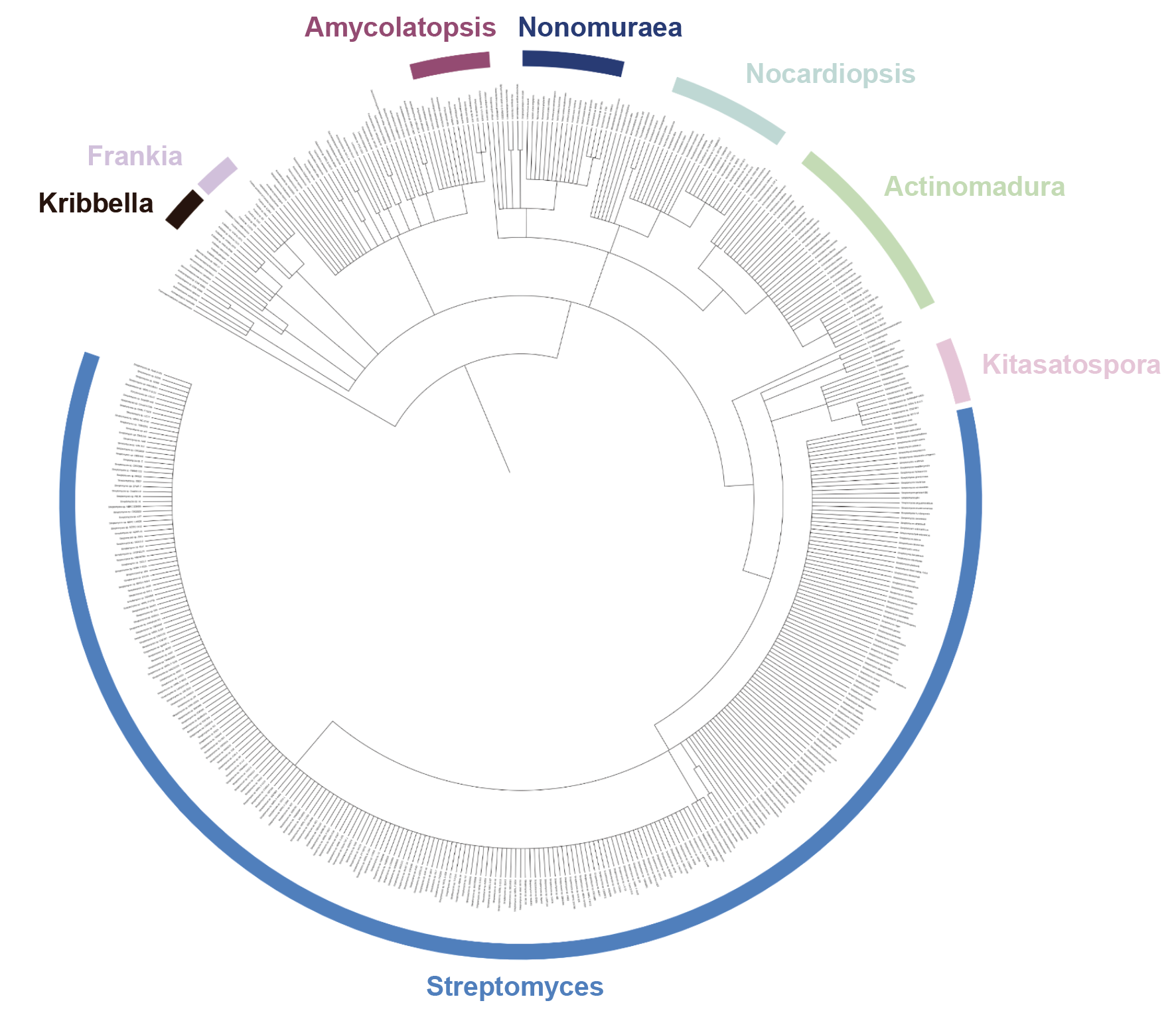

**Figure S4: Phylogenetic Trees of Organisms that Encode Imiditide BGCs.** All imiditide BGCs are encoded in Gram-positive bacterial genomes. The most representative genus that encode imiditide BGCs include *Streptomyces*, *Actinomadura*, *Nocardiopsis*, *Nonomuraea*, and *Amycolatopsis*.

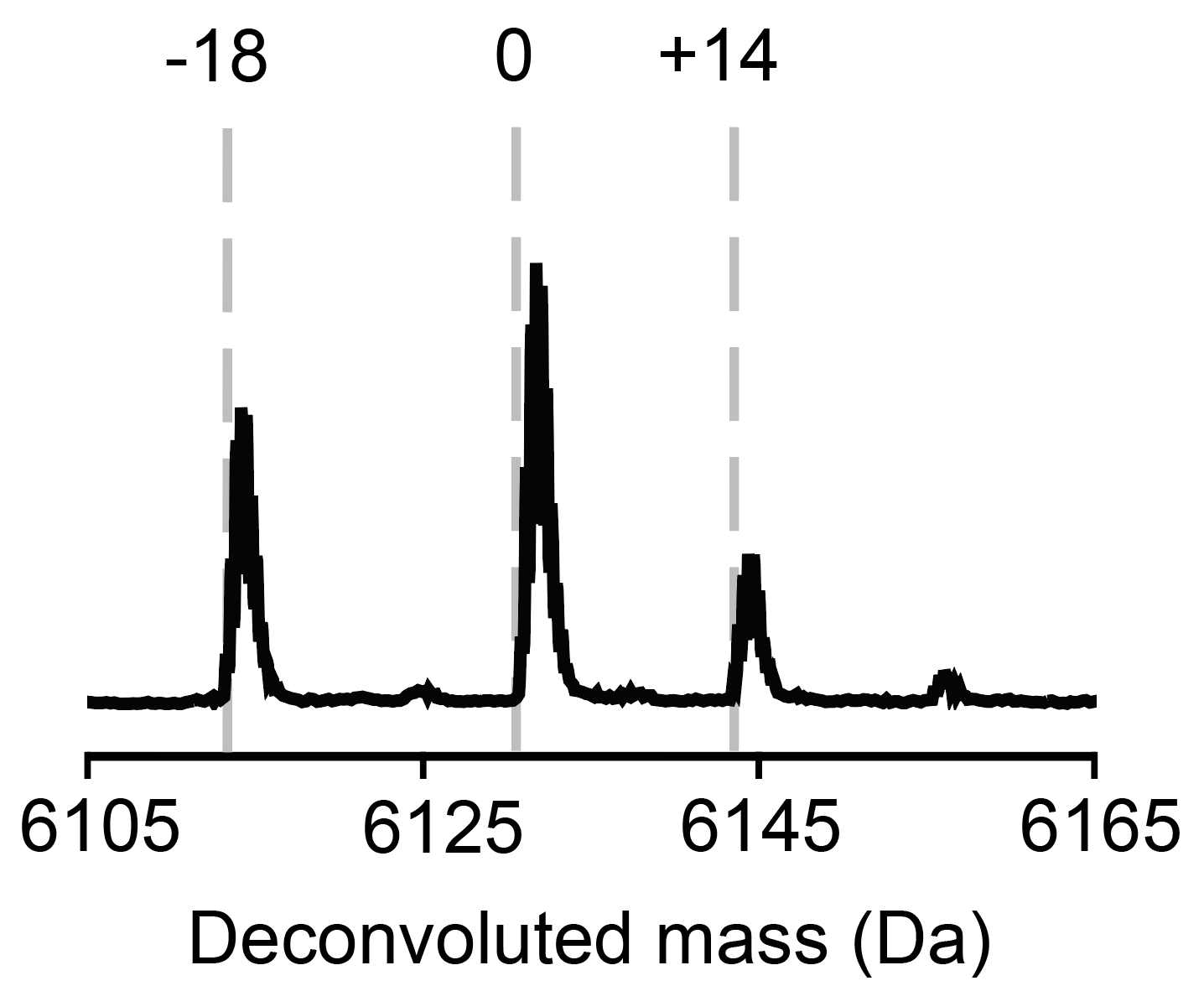

**Figure S5: Coexpression of His_6_-NmaA and NmaM.** Mass spectrum of modified His6-NmaA shows the emergence of the putative methylated and aspartimidylated species, showing that SUMO does not affect the post-translational modification. However, since the SUMO tag greatly enhanced the level of NmaA expression, we used His_6_-SUMO-NmaA for all subsequent heterologous expressions.

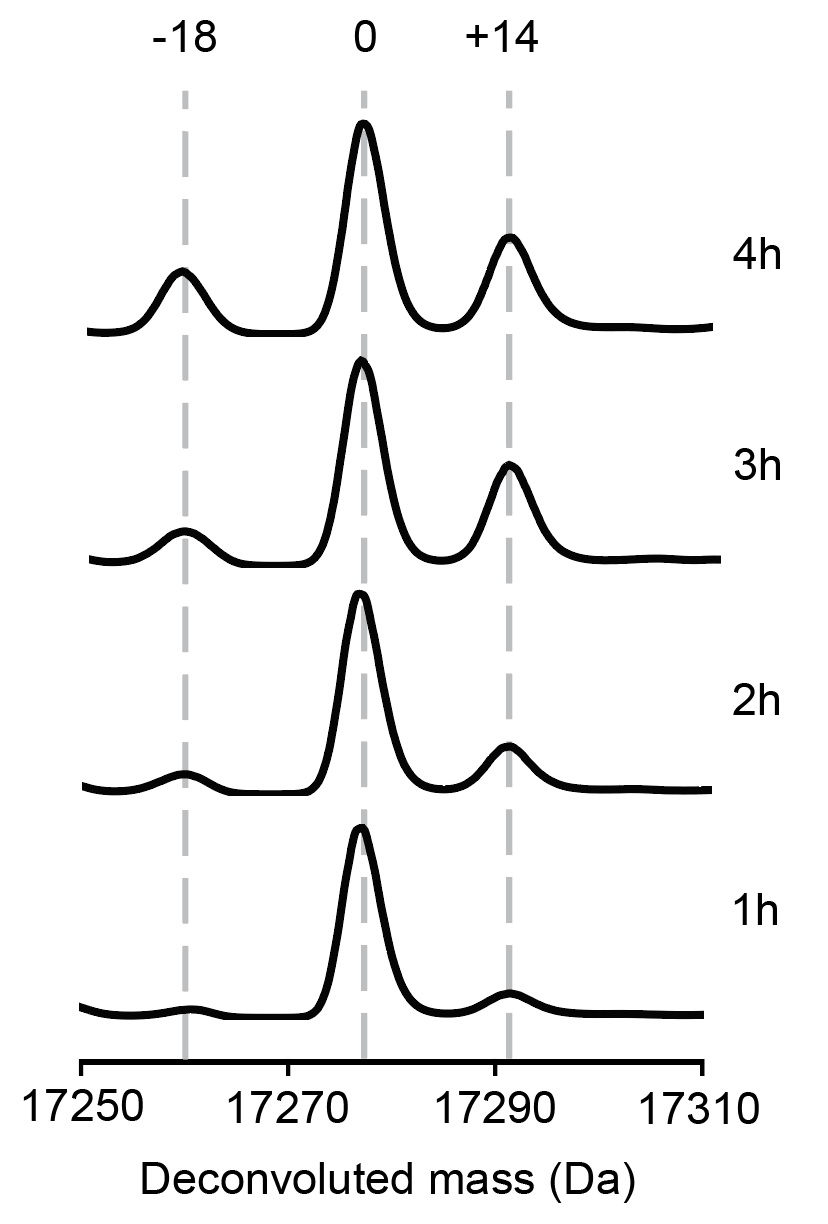

**Figure S6: Time Course of Heterologous Expressions of mNmaA^M^ in *E. coli* BL21 (DE3) *ΔslyD***. Mass spectra of heterologous expressions of mNmaA^M^ for 1, 2, 3, and 4 hours are shown here. The methylation step occurred first and accumulated. Then the methylated mNmaA formed an intramolecular aspartimide spontaneously.

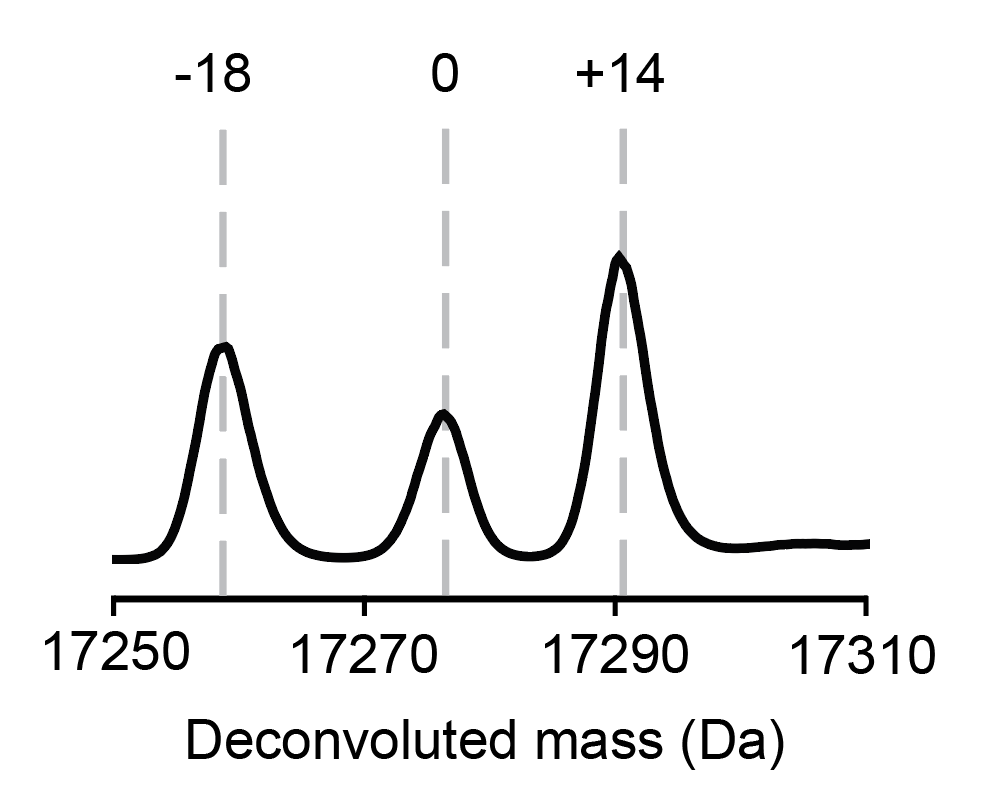

**Figure S7: Heterologous Expression of mNmaA^M^ using *E. coli* BL21 (DE3) *ΔpcmΔslyD* strain for 20h.** The extent of methylation and aspartimidylation observed at 20h of expression was similar compared to heterologous expression of mNmaA^M^ using *E. coli* BL21 (DE3) *ΔslyD* strain. Therefore, all subsequent heterologous expression experiments were carried out using *E. coli* BL21 (DE3) *ΔslyD* strain.

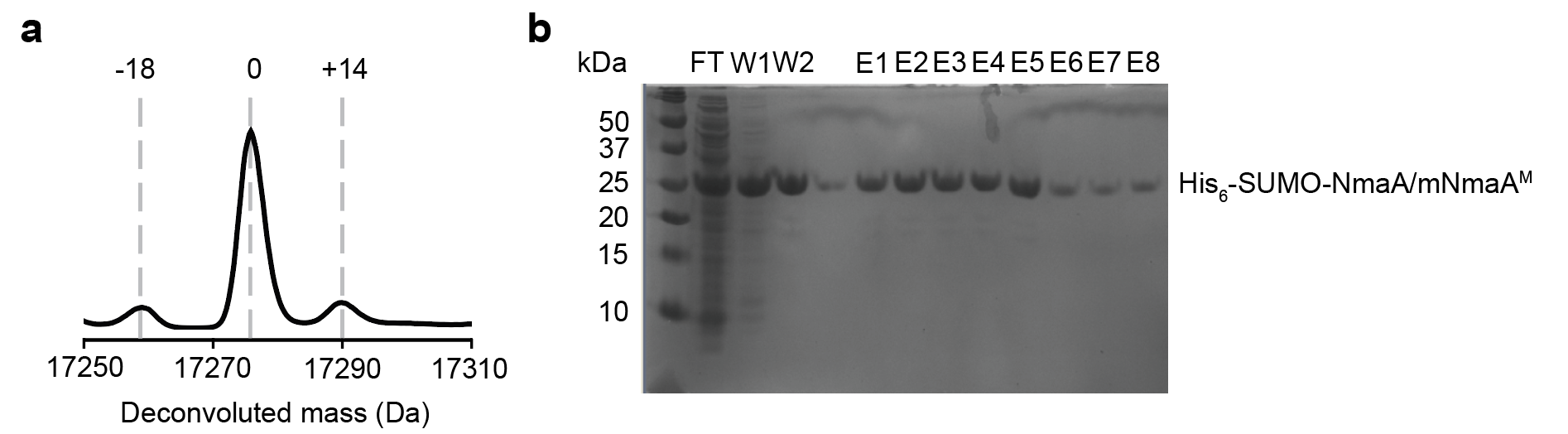

**Figure S8: Heterologous Expression of mNmaA^M^ with NmaM Constitutively Expressed. a)** Mass spectrum of mNmaA^M^ when NmaM was expressed constitutively. Much lower levels of methylated and aspartimidylated species were detected, showing that the extent of modification was proportional to the level of NmaM produced. **b)** SDS-PAGE gel of mNmaA^M^ showed no pulldown of NmaM, suggesting that the expression level of NmaM was low. FT: Flowthrough. W: Wash. E: Elution.

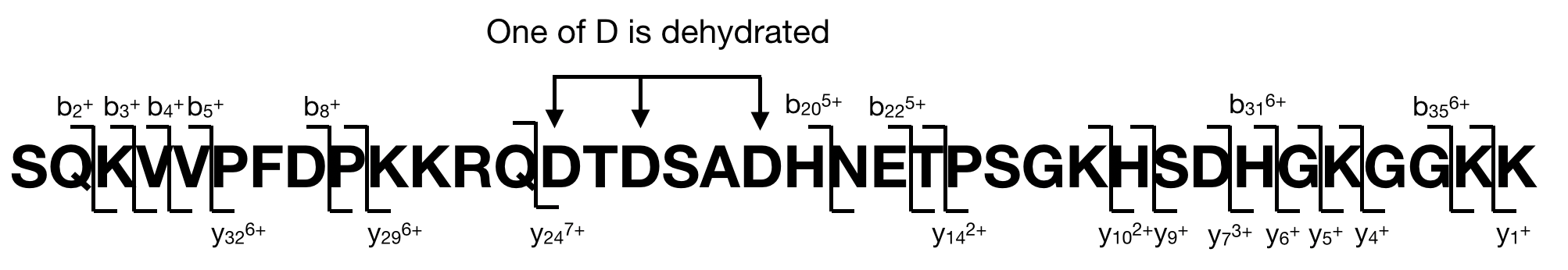

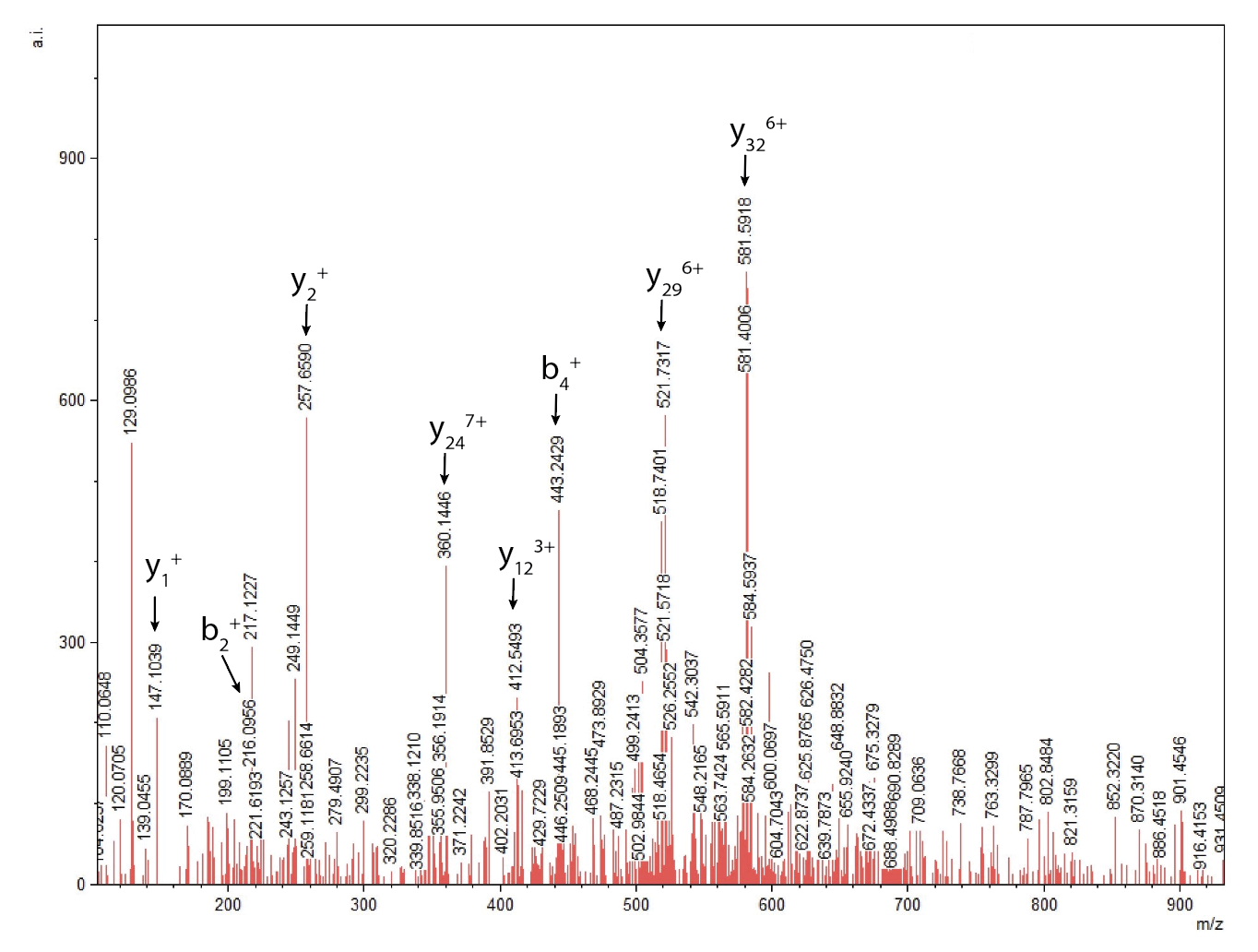

| **Detected fragment** | **Expected mass** | **Observed**  **mass** |
| --- | --- | --- |
| b_2_^+^ | 216.0979 | 216.0956 |
| b_3_^+^ | 344.1928 | 344.1780 |
| b_4_^+^ | 443.2613 | 443.2429 |
| b_5_^+^ | 542.3297 | 542.3037 |
| b_8_^+^ | 901.4778 | 901.4546 |
| b_20_^5+^ | 453.2311 | 453.2266 |
| b_22_^5+^ | 501.8482 | 501.8683 |
| b_31_^6+^ | 576.1123 | 576.1652 |
| b_35_^6+^ | 585.6159 | 585.6056 |
| y_1_^+^ | 147.1128 | 147.1039 |
| y_4_^+^ | 389.2507 | 389.2423 |
| y_5_^+^ | 517.3457 | 517.3504 |
| y_6_^+^ | 574.3671 | 574.3522 |
| y_7_^3+^ | 237.8135 | 237.8117 |
| y_9_^+^ | 913.4850 | 913.4775 |
| y_10_^2+^ | 525.7756 | 525.7525 |
| y_14_^2+^ | 710.3762 | 710.3865 |
| y_24_^7+^ | 356.1669 | 356.1914 |
| y_29_^6+^ | 521.5938 | 521.5718 |
| y_32_^6+^ | 581.4519 | 581.4006 |

**Figure S9: MS/MS Fragmentation of NmaA^M^ (10-46).** Tables of expected and detected masses of fragments are provided. The fragmentation pattern indicated that one of the D23, D25, and D28 was dehydrated.

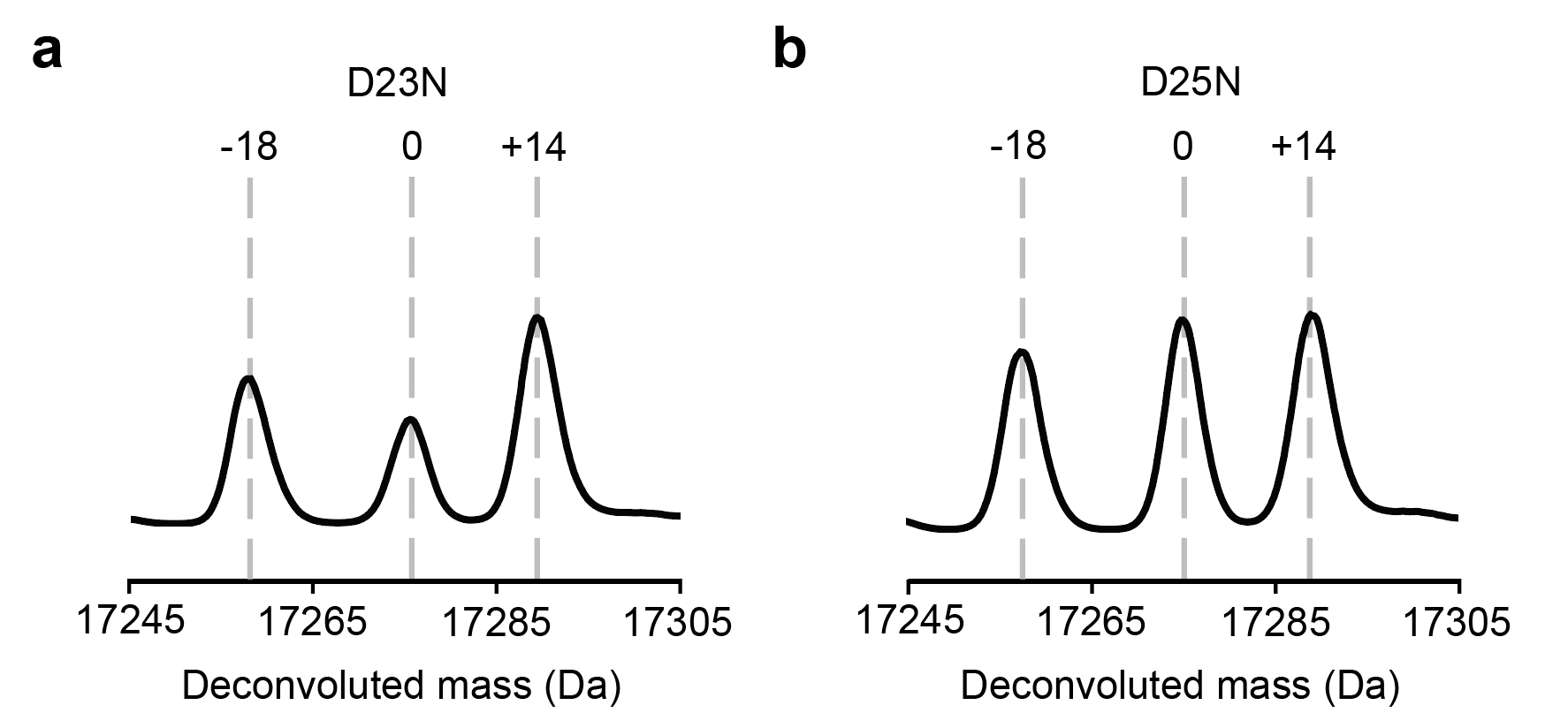

**Figure S10: D23 and D25 are not the Sites of Aspartimidylation. a)** Mass spectrum of His_6_-SUMO-NmaA D23N variant coexpressed with NmaM. Methylation and aspartimidylation still occurred, showing that D23 was not involved in the aspartimide formation. **b)** Mass spectrum of His_6_-SUMO-NmaA D25N variant coexpressed with NmaM. Similar to D23N variant, the D25N variant could still be methylated and aspartimidylated, showing that D25N variant was not involved in the aspartimide formation.

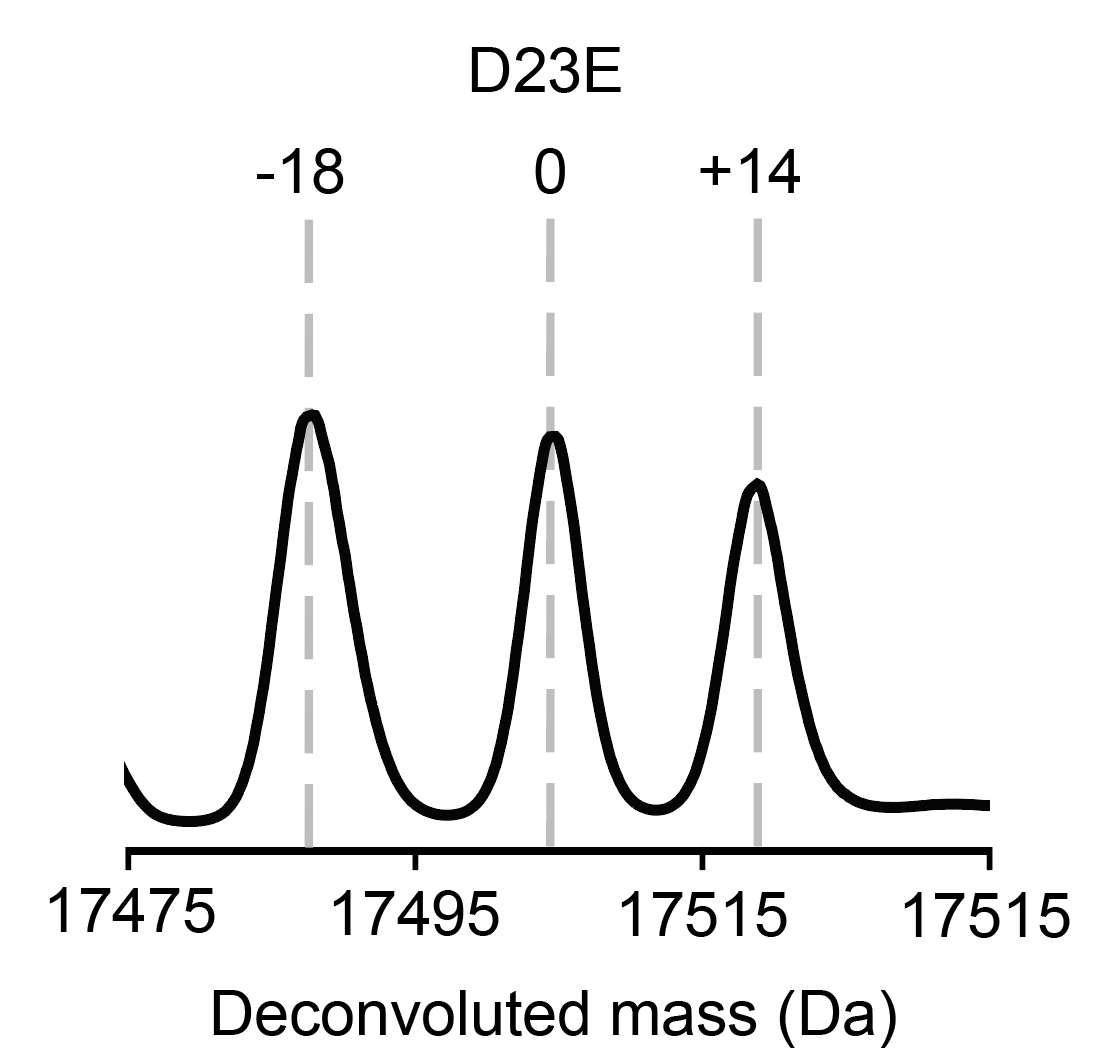

**Figure S11: His_6_-SUMO-NmaA D23E Variant can be Modified as Efficiently as the Wildtype His_6_-SUMO-NmaA.** Mass spectrum of mNmaA^M^ D23E. The aspartimide can form as efficiently as wildtype. The mass contains the first methionine in the protein sequence (start with MRGSHHHHHHGS) compared to other constructs (start with MSGSHHHHHHGS). mNmaA^M^ D23E was further digested with endopeptidase GluC to generate a short peptide fragment for FTIR measurements.

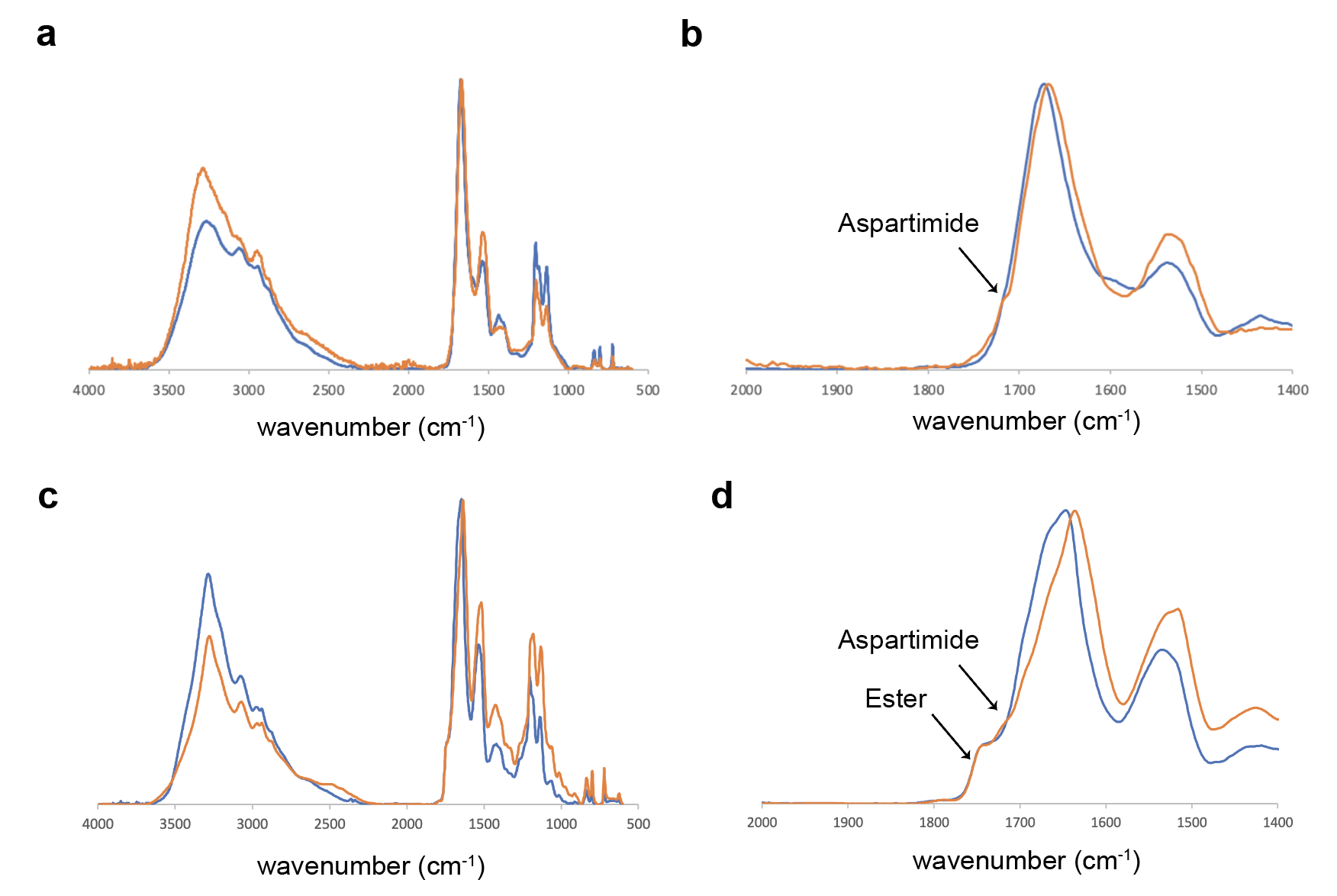

**Figure S12: FTIR Spectrum of mNmaA^M^ (24-46) Confirmed mNmaA^M^ Contains an Aspartimide. a)** Overlay of FTIR spectra of mNmaA^M^ (24-46) (orange) and unmodified NmaA (24-46) (blue). **b)** As in **a)**, but zoomed in the amide signal region. NmaA^M^ (24-46) showed an additional shoulder peak at 1708 cm^-1^. **c)** Overlay of FTIR spectra of amycolimiditide (orange) and pre-amycolimiditide (blue). **d)** As in **c)**, but zoomed in the amide signal region. Both amycolimiditide and pre-amycolimiditide showed a peak at 1742 cm^-1^ signaling the formation of esters. An additional shoulder peak at 1708 cm^-1^ presented for the amycolimiditide sample, suggesting that it corresponded to the aspartimide formation.

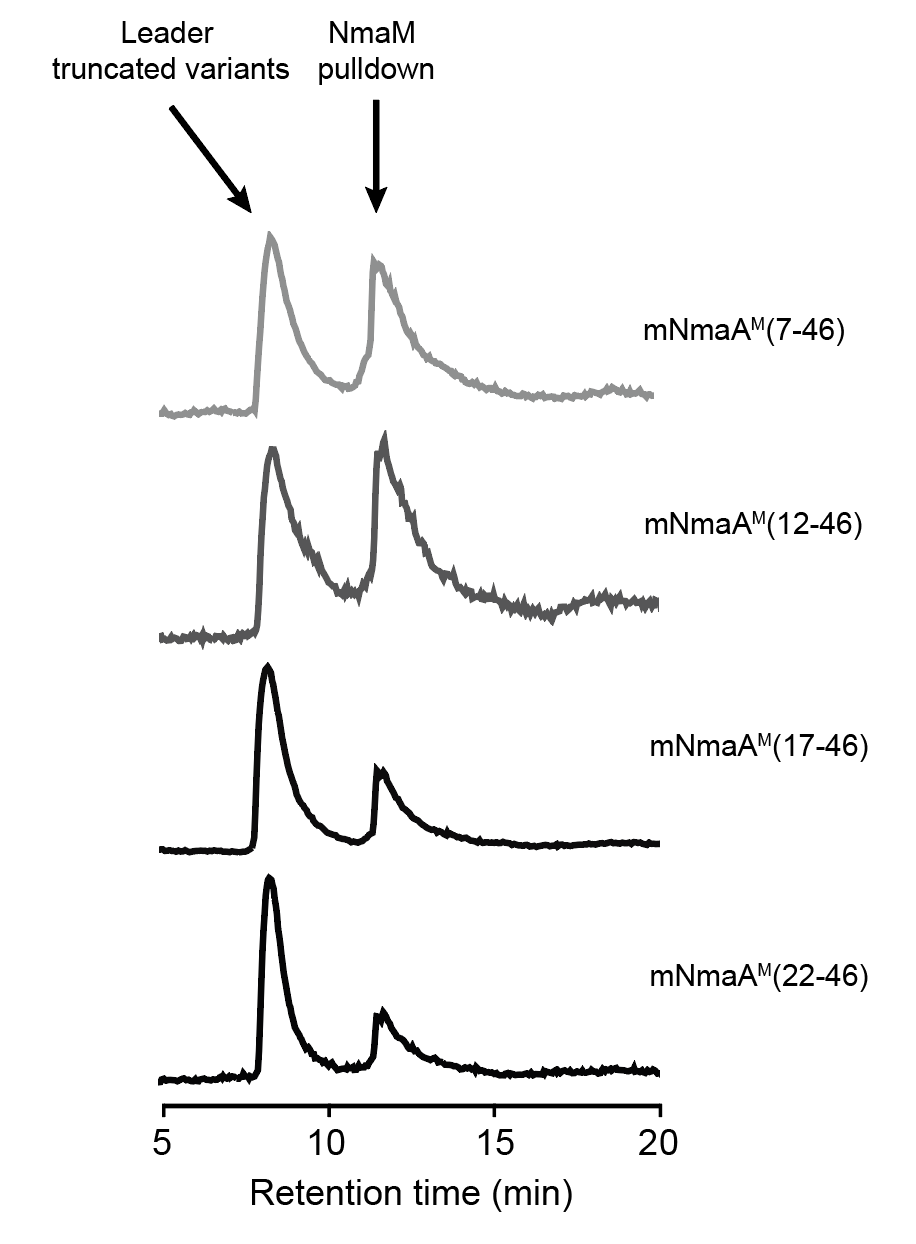

**Figure S13: Total Ion Current Chromatograms (TICs) of mNmaA^M^ Leader Truncated Variants.** The leader truncated variants eluted around 8 mins, while the untagged NmaM that got pulled down eluted at 12 mins. The removal of NmaA (12-21), which had a high content of basic amino acids, reduced the amount of NmaM that mNmaA^M^ could pull down, showing this region was crucial for substrate recognition.

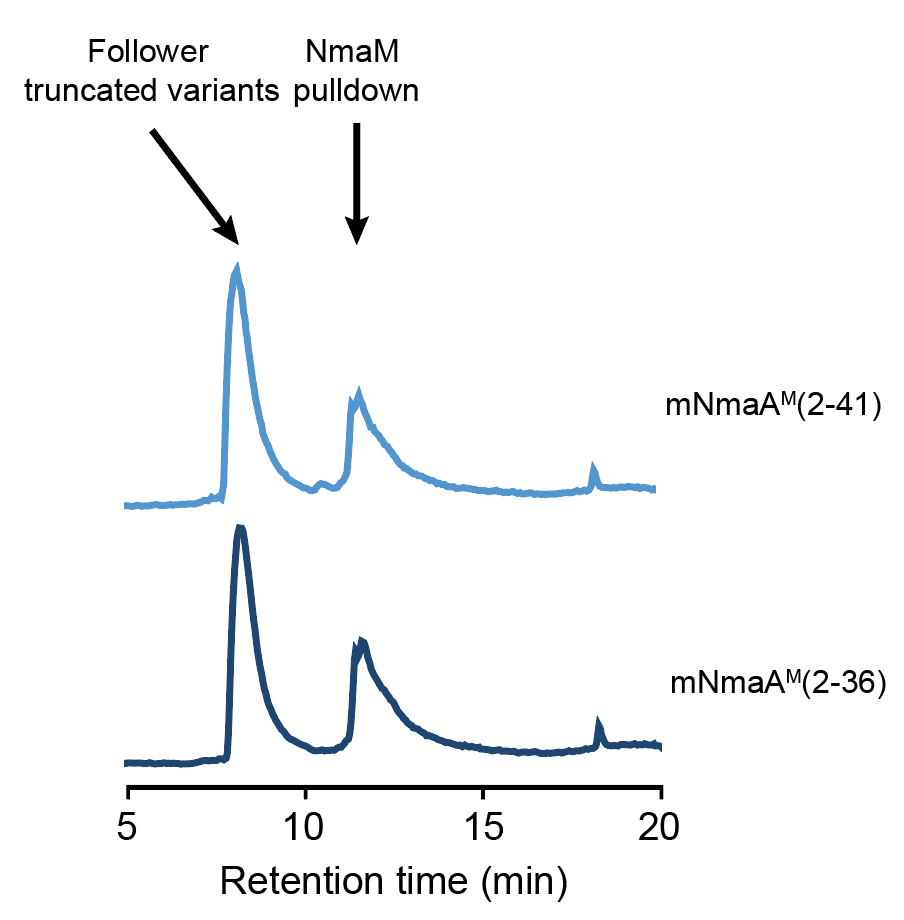

**Figure S14: Total Ion Current Chromatograms (TICs) of mNmaA^M^ Follower Truncated Variants.** The follower truncated variants eluted at 8 mins, while the untagged NmaM that got pulled down eluted at 12 mins. The removal of NmaA (42-46), which were rich in Lys, reduced the amount of NmaM that mNmaA^M^ could pull down, showing this region was crucial for substrate recognition.

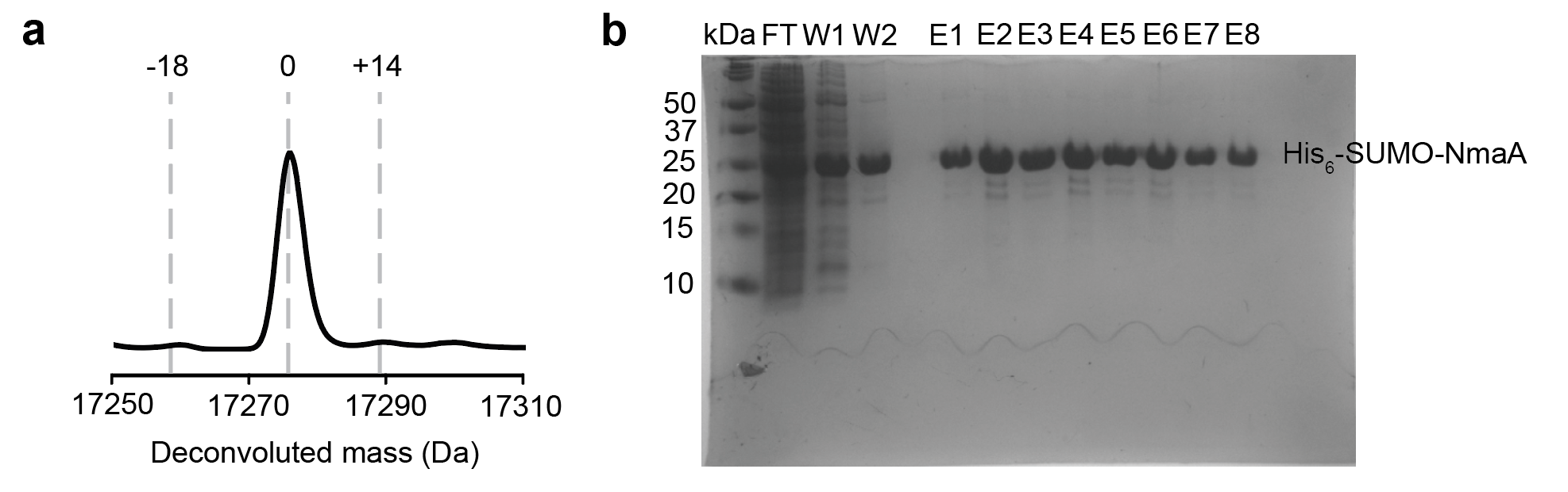

**Figure S15: Heterologous Coexpression of His_6_-SUMO-NmaA and NmaMΔC50. a)** No modification was observed on the precursor, suggesting that the C-terminal part that contained the 41 aa motif for bioinformatic study was crucial for substrate recognition and subsequent aspartimide formation. **b)** SDS-PAGE gel of purified His_6_-SUMO-NmaA fractions show the precursor did not pull down NmaMΔC50.

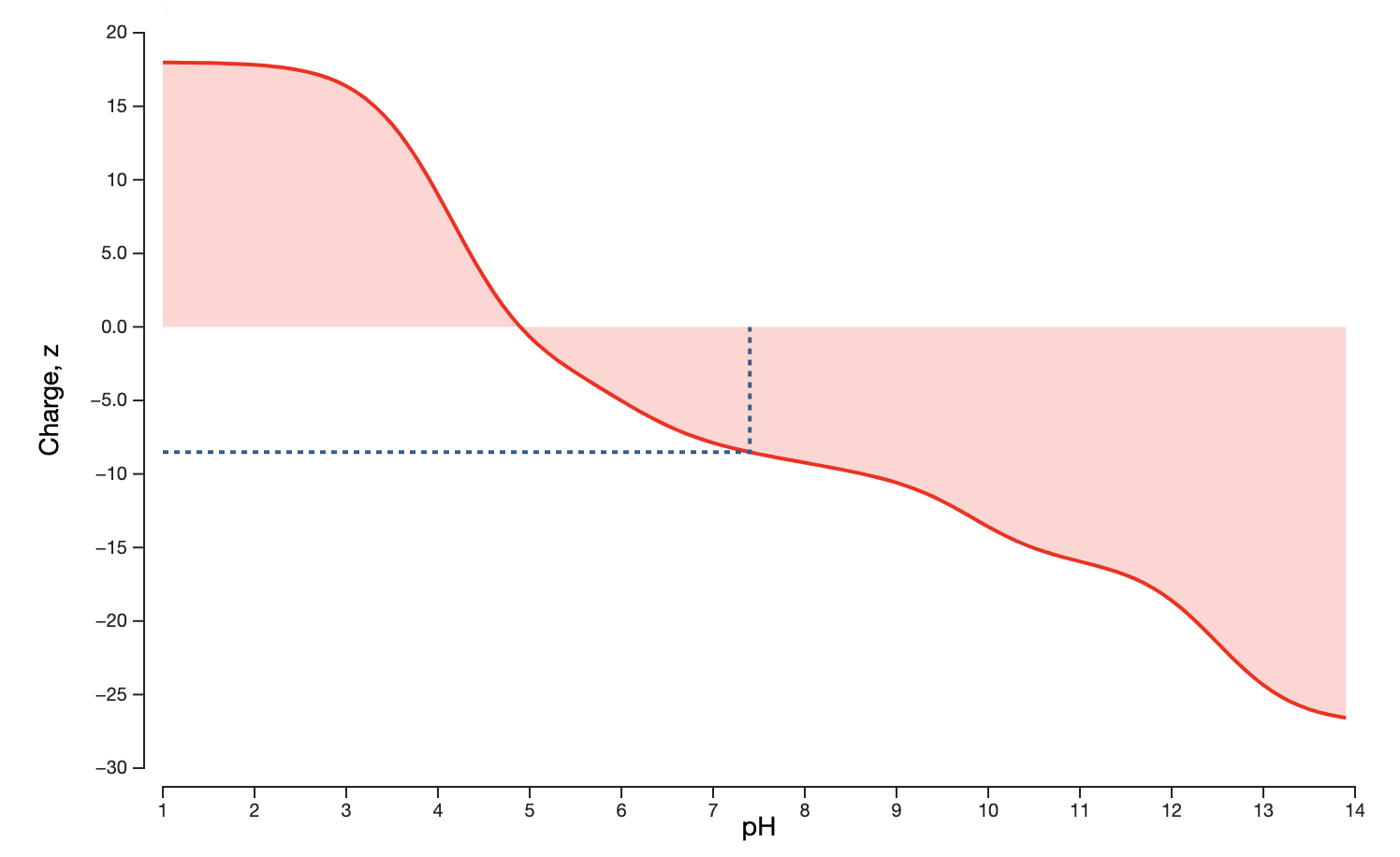

**Figure S16: Titration Curve of the C-terminal Domain of NmaM.** The C-terminal domain of NmaM has a pI value of 4.90, showing that it contains a high content of residues with acidic side chains. At physiological pH of 7.4, the C-terminal domain of NmaM contains a net charge of -8.5 (shown in blue dotted line).

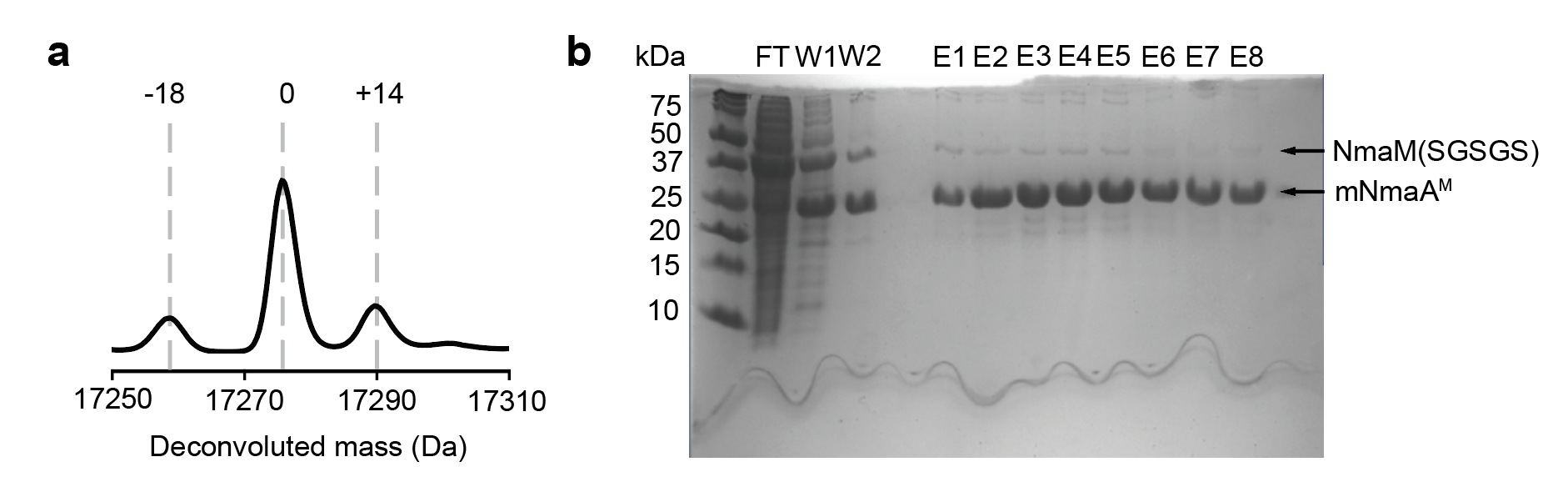

**Figure S17: Heterologous Expression of mNmaA^M^** **Using NmaM (SGSGS). a)** Mass spectra of mNmaA^M^ modified by NmaM (SGSGS). Removing 4 negative charges are in the C-terminal domain of NmaM was detrimental to the extent of modification on the precursor, showing that this these negative charges were important for the post-translational modification. **b)** SDS-PAGE gel of purified mNmaA^M^ showed that mNmaA^M^ pulled down less NmaM(SGSGS) compared to wildtype NmaM, indicating that the charge-charge interactions between NmaA and NmaM are important for substrate recognition.

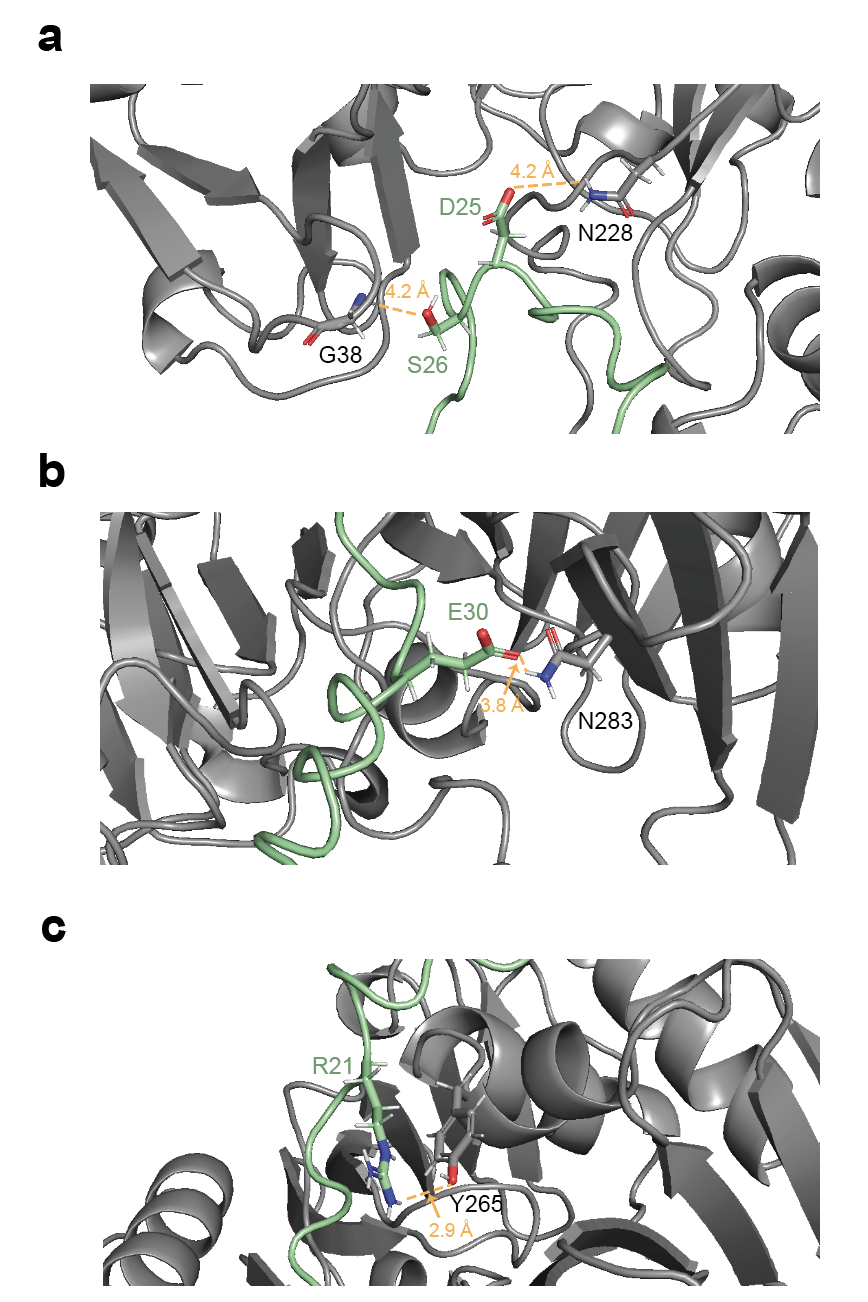

**Figure S18: Charge-charge and Hydrogen Bond Interactions Between NmaA and NmaM Predicted by AlphaFold.** NmaA and NmaM are colored palegreen and grey, respectively. Core region of NmaA interacts extensively with NmaM. **a)** D25 and S26 in NmaA interact with N228 and G38 in NmaM, respectively. **b)** E30 in NmaA interacts with N283 in NmaM. **c)** R21 in NmaA interacts with Y265 in NmaM.

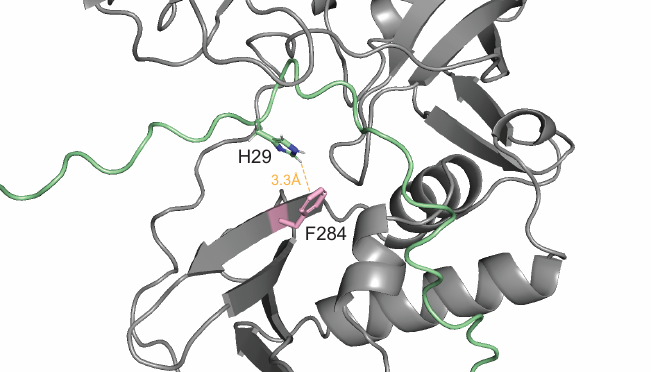

**Figure S19: AlphaFold Predicts Interactions between the *n* + 1 Residue H29 in NmaA and F284 in NmaM.** NmaA and NmaM are colored palegreen and grey, respectively. F284 residue in NmaM is shown in pink. Both sidechains of H29 in NmaA and F284 in NmaM are shown. The H in the imidazole ring in the sidechain of H29 is predicted to be 3.3 Å away from the π system in the aromatic ring in the sidechain of F284. This interaction can explain why H29 is the preferred *n* + 1 residue for mNmaA^M^.

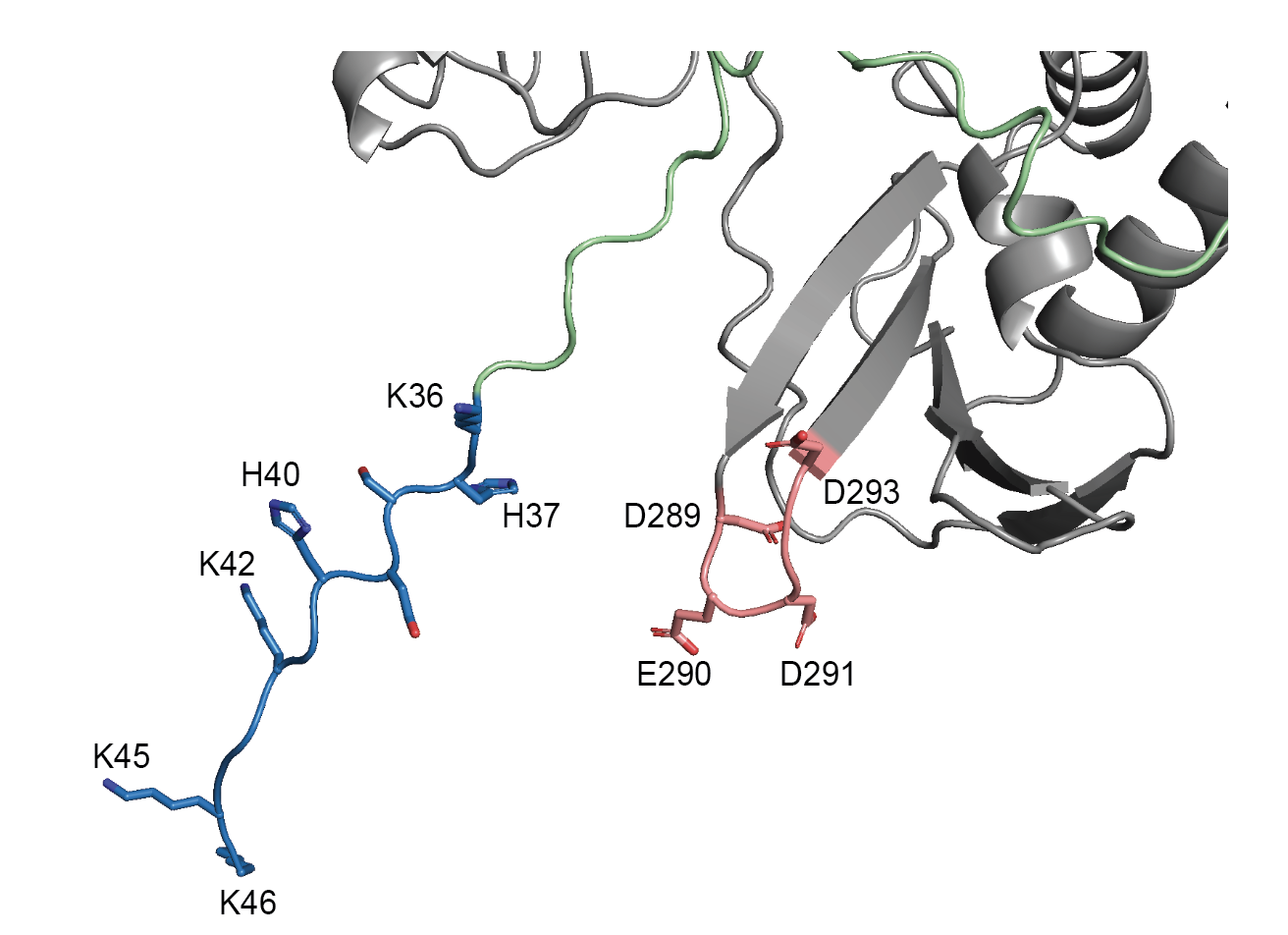

**Figure S20: The Acidic Loop in NmaM Should be Engaged with the Follower Sequence.** NmaA and NmaM are colored palegreen and grey, respectively. The acidic loop in NmaM is colored pink, and the follower of NmaA is colored blue. Sidechains of acidic residues in the acidic loop in NmaM and the basic residues in the follower sequence in NmaA are shown. Alphafold model likely misses the interactions between these two parts as our experimental results suggest that substitutions of each part results in less interactions between NmaA and NmaM to similar degrees.

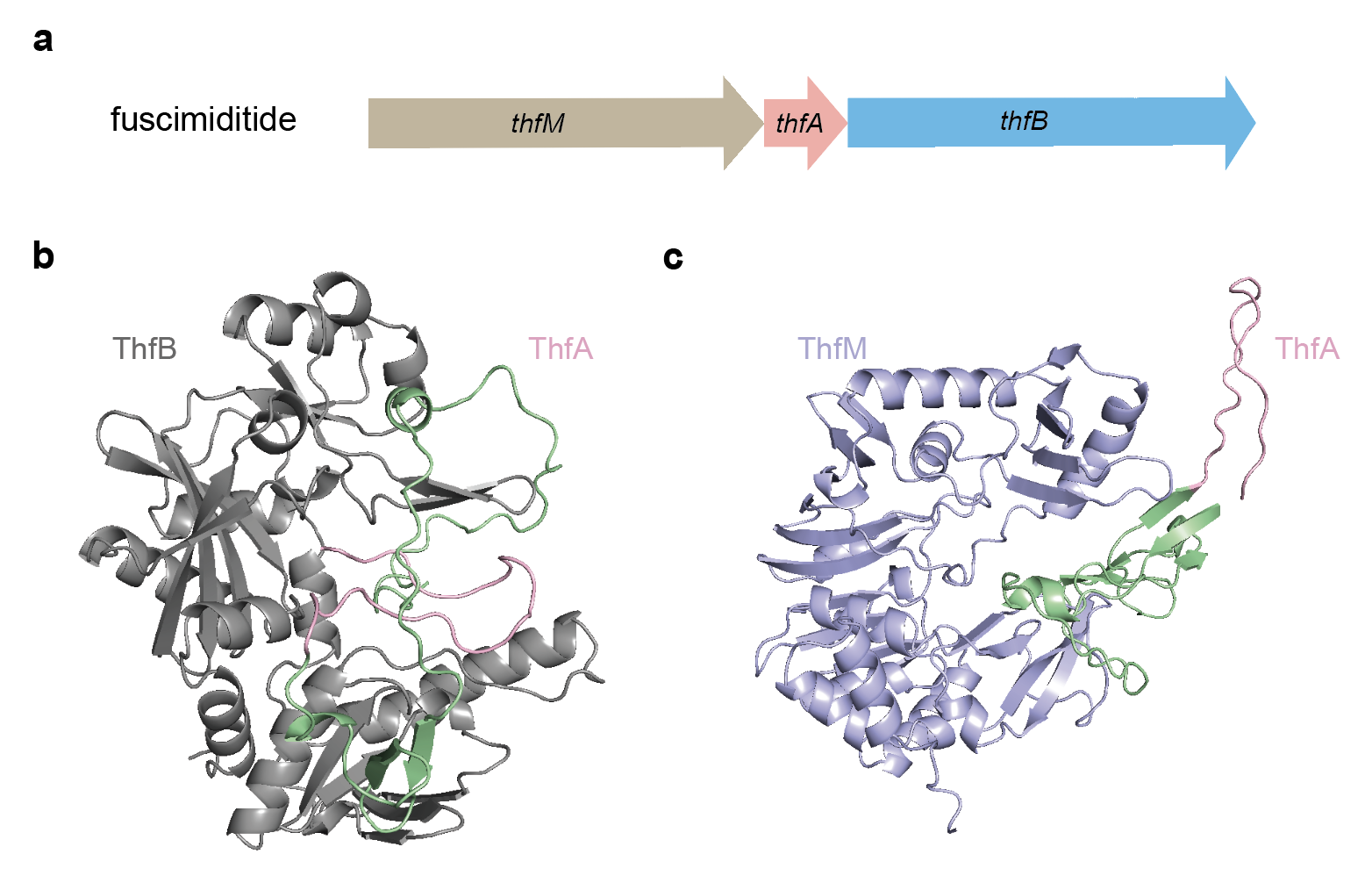

**Figure S21: Alphafold Predicts the Order of Modifications in RiPP Biosynthetic Clusters. a)** Fuscimiditide BGC.^5^ *thfA* encodes the precursor, *thfB* encodes an ester-installing ATP-grasp enzyme, and thfM encodes an *O*-methyltransferase that installs an aspartimide. In fuscimiditide biosynthesis, 2 ester linkages form first by ThfB and an aspartimide is installed after by ThfM. The reaction order is strict. **b)** Alphafold predicts that ThfB reacts with ThfA. ThfB is colored grey. The leader and core of ThfA are colored palegreen and pink, respectively. **c)** Alphafold predicts that ThfM does not react with ThfA. ThfA adapts the same color scheme as in **b)**, and ThfM is colored in lightblue.

### Supplementary Tables

Table S1: A list of 50 *O*-methyltransferase NCBI Accession Numbers in Putative Novel RiPP Family BGC by a BLASTP on *O*-methyltransferase from *Nonomuraea jiangxiensis* found in ThfM SSN

| Organism | NCBI Accession Number |
| --- | --- |
| *Nonomuraea jiangxiensis* | WP_090935243.1 |
| *Nonomuraea candida* | WP_052423694.1 |
| *Nonomuraea candida* | WP_043634548.1 |
| *Nonomuraea candida* | WP_052422810.1 |
| *Nonomuraea maritima* | WP_090766369.1 |
| *Nonomuraea sp. terrae* | WP_132615232.1 |
| *Nonomuraea aridisoli* | WP_235854837.1 |
| *Nonomuraea cypriaca* | WP_195896576.1 |
| *Nonomuraea cypriaca* | WP_195900101.1 |
| *Nonomuraea sp. ATCC 55076* | WP_186404513.1 |
| *Sphaerisporangium rosea* | WP_113978463.1 |
| *Sphaerisporangium rosea* | WP_113977729.1 |
| *Sphaerisporangium rosea* | WP_225878617.1 |
| *Sphaerisporangium rosea* | WP_158578444.1 |
| *Desertiactinospora gelatinilytica* | WP_111169254.1 |
| *Desertiactinospora gelatinilytica* | WP_158557932.1 |
| *Desertiactinospora gelatinilytica* | WP_111172032.1 |
| *Desertiactinospora gelatinilytica* | WP_111171751.1 |
| *Nonomuraea sp. C10* | WP_261808771.1 |
| *Nonomuraea phyllanthi* | WP_228010834.1 |
| *Nonomuraea deserti* | WP_165974922.1 |
| *Nonomuraea jabiensis* | WP_246556574.1 |
| *Nonomuraea diastatica strain KC712* | WP_165977030.1 |
| *Nonomuraea mesophile strain 6K102* | WP_132637211.1 |
| *Nonomuraea lactucae strain NEAU-YG30 C5429* | WP_113704943.1 |
| *Nonomuraea wenchangensis* strain CGMCC 4.5598 | WP_091090976.1 |
| *Nonomuraea sp. PA05* | WP_148441289.1 |
| *Nonomuraea sp. PA05* | WP_148434645.1 |
| *Nonomuraea phyllanthi* | WP_193319035.1 |
| *Nonomuraea coxensis DSM 45129* | WP_020542946.1 |
| *Sinosporangium album* | WP_245691186.1 |
| *Streptosporangium violaceochromogenes* | GHE42692.1 |
| *Thermocatellispora tengchongensis* | WP_185051145.1 |
| *Thermocatellispora tengchongensis* | WP_185049748.1 |
| *Thermomonospora echinospora* | WP_235018084.1 |
| *Thermomonospora amylolytica* | WP_119728020.1 |
| *Thermomonospora cellulosilytica* | WP_182703951.1 |
| *Actinomadura craniellae* | WP_111866096.1 |
| *Actinomadura craniellae* | WP_111870683.1 |
| *Actinomadura oligospora* | WP_051467946.1 |
| *Actinomadura hallensis* | WP_246077203.1 |
| *Actinomadura madurae* | WP_111831063.1 |
| *Actinomadura sp. NAK00032* | WP_254715715.1 |
| *Actinomadura latina* | WP_067627936.1 |
| *Actinomadura darangshiensis* | WP_132199480.1 |
| *Actinomadura sp. 7K507* | WP_165966849.1 |
| *Actinomadura sp. 6K520* | WP_131980484.1 |
| *Nocardiopsis dassonvillei* | WP_174546145.1 |
| *Nocardiopsis alborubida* | WP_061082159.1 |
| *Actinomadura oligospora strain ATCC 43269* | WP_051467946.1 |

Table S2: Primers and constructs used in this study.

| Primer | Primer Sequence | Primer Description | Plasmid construct | Plasmid Description |
| --- | --- | --- | --- | --- |
| LC49 | GCATCCATGGATGGGCCGTCATGCGGAAGATCCGGAAAG | Primer-1-NmaA | pRSFDuet | pLC52 |
| LC50 | TATCCTGACGTTTTTTCGGATCAAACGGCACCACTTTCTGGCTTTCCGGATCTTCCGCAT | Primer-2-NmaA | pRSFDuet | pLC52 |
| LC51 | GATCCGAAAAAACGTCAGGATACCGATAGCGCGGATCATAATGAAACCCCGAGCGGGAAA | Primer-3-NmaA | pRSFDuet | pLC52 |
| LC52 | GTACAAGCTTTCATTTTTTGCCGCCTTTGCCATGATCGCTATGTTTCCCGCTCGGGGTTT | Primer-4-NmaA | pRSFDuet | pLC52 |
| LC53 | TCACACAGAATTCATTAAAGAGGAGAAATTAACTAT | pQE80-EcoRI forward | pQE80 | pLC53, 71-73, 80-83, 86, 88-91 |
| LC54 | GCATGGTCTCTACCAATCTGTTCTCTGTGAG | pQE80-SUMO reverse (for SUMO-NmaA) | pQE80 | pLC53 |
| LC55 | gcatGGTCTCctggtTCCGGCCGTCATGCGGAAGAT | pQE80-NmaA forward(for SUMO-NmaA) | pQE80 | pLC53 |
| LC56 | gcatGGTCTCTagctttcattttttgccgccttt | pQE80-NmaA hindIII reverse(for SUMO-NmaA) | pQE80 | pLC53, 59, 71-73,80-83, 86, 90-91 |
| LC57 | gtaccatatgCTGAGCCTGAGCG | NmaM NcoI forward | pRSFDuet | pLC54, 55, 92, 95 |
| LC58 | gcatcctaggTCACGGCAACG | NmaM AvrII reverse | pRSFDuet | pLC54, 55, 92 |
| LC59 | GCACGGATCCGGCCGTCATGCGGAAGAT | His_6_-NmaA BamHI forward | pQE80 | pLC59 |
| LC60 | AAAACGTCAGAATACCGATAGCG | D23N internal forward | pQE80 | pLC71 |
| LC61 | CGCTATCGGTATTCTGACGTTTT | D23N internal reverse | pQE80 | pLC71 |
| LC62 | GTCAGGATACCAATAGCGCG | D25N internal forward | pQE80 | pLC72 |
| LC63 | TTTCATTATGATtCGCGCTATCGG | D25N internal reverse | pQE80 | pLC72 |
| LC64 | CCGATAGCGCGaATCATAATGAAA | D28N internal forward | pQE80 | pLC73 |
| LC65 | TTTCATTATGATtCGCGCTATCGG | D28N internal reverse | pQE80 | pLC73 |
| LC66 | GCATGGTCTCTACCAATCTGTTCTCTGTGAG | Leader truncation SUMO reverse | pQE80 | pLC80-83 |
| LC67 | gcatGGTCTCctggtTCCGATCCGGAAAGCCAGAAAGT | Leader -5 aa BsaI forward | pQE80 | pLC80 |
| LC68 | gcatggtctcctggttccAAAGTGGTGCCGTTTGATC | Leader -10 aa BsaI forward | pQE80 | pLC81 |
| LC69 | gcatggtctcctggttccgatccgaaaaaacgtcaggatacc | Leader -15 aa BsaI forward | pQE80 | pLC82 |
| LC70 | gcatggtctcctggttccCAGGATACCGATAGCGCG | Leader -20 aa BsaI forward | pQE80 | pLC83 |
| LC71 | AAAACGTCAGGAaACCGATAGCG | D23E internal forward | pQE80 | pLC86 |
| LC72 | CGCTATCGGTtTCCTGACGTTTT | D23E internal reverse | pQE80 | pLC86 |
| LC73 | gcatGGTCTCTagctttcagccatgatcgctatgtttcc | follower -5 aa BsaI reverse | pQE80 | pLC88 |
| LC74 | gcatGGTCTCTagctttcatttcccgctcggggtt | follower -10 aa BsaI reverse | pQE80 | pLC89 |
| LC75 | TAGCGCGGATGGTAATGAAACC | H29G forward | pQE80 | pLC90 |
| LC76 | GGTTTCATTACCATCCGCGCTA | H29G reverse | pQE80 | pLC90 |
| LC77 | TAGCGCGGATACTAATGAAACC | H29T forward | pQE80 | pLC91 |
| LC78 | GGTTTCATTAGTATCCGCGCTA | H29T reverse | pQE80 | pLC91 |
| LC79 | AGTGGCAGCGGTAGTTGCTTTCGTCTGTGGATTAGC | NmaM SGSGS forward | pRSFDuet | pLC92 |
| LC80 | AACTACCGCTGCCACTGGTTTCCGCGCTAAAATTGC | NmaM SGSGS reverse | pRSFDuet | pLC92 |
| LC81 | gcatcctaggtcaATACACTTCATAATCTTTACGATTGGGACG | NmaM ΔC50aa reverse | pRSFDuet | pLC95 |
| LC82 | TCCAAGCTAGCcatcaattaagaaaaa | mcjBCD promoter NheI Forward | pQE80 | pLC99 |
| LC83 | AGGCTCAGCATATGTATATCTCCTTCTTATACTTA | mccjBCDNmaM internal reverse | pQE80 | pLC99 |
| LC84 | GGAGTGTGCaAaATGCTGAGCCTGAGCGAT | MccjBCDNmaM internal forward | pQE80 | pLC99 |
| LC85 | gcatccatggTCACGGCAACGGATGGTC | NmaM NcoI reverse | pQE80 | pLC99 |

Table S2. Sequence of gblock for construction of NmaM

| gblock | Sequence |
| --- | --- |
| 1 | TTTAAcatATGCTGAGCCTGAGCGATCTGCCGGAAAATACGCCCGACGAAATCCGTAGCGCGATTCGTGCGGTGCCGCGTCATCAGTTTATTCCGGTGGTGGCGTTAGTGACTCCCCCTGGTACGGCTCCGCGTCTGATTGATCGTGATGCAGATCCGCGTGCTTGGTGGGATGCGGTGTATAGCAATACCCCGGTTATTACCCAACTGAATGATGGCGCGACCCCAGTGCGTGATCTGATAGGCCGTTATACTAGCAGCGCGAGCGCGCCTAGCACCGTGGCTGATATGCTGACCCTGTTGCGTCCTGCGCCTGGTCATCGTGTTCTGGAAATTGGCACTGGCACCGGCTGGACGAGCGCTTTACTGTGCCATCTGGTTGGGGGTCGTGGCCAGGTGACGAGCATTGAAATAGATCAAACGGTGGCGGAACAGGCGGCGAAAAATCTGGCGGGAGCAGGAGCGCCGGTGACTTTAGTCGTGGGCGACGGCGCGTTGGGATGGCCGCCAGGGGAACCGTATGATCGGGTTCATGTGACCTGCGGCGTGCGTACCGTGCCGTATGCGTGGGTCGAACAGTGTCGTCCTGGTGCAGTCATTGCGCTGCCTTATTGCCCCGATTTTGGCGGAGGCCATAGCCTGAAACTGGTCGTGCGTCCGGATGGCGTGGCGATCGGCCGTTTCCCGGGCAATGCCTCCTACATGATGAGCCGTCCGCAGCGGCCACGTCCGACCCTGGAAGCGCGTGGCCCGCATGAGCAGCGTTCCGCGACCACTCGTGTGGACCCGCGTACCATTGCGTATGCTCCGCCTGGCGCGGATTTGGCCATAAGCGCATTAACCGGCCTGGTGAGCAATTTTAGCGCGGAAACCGATGAAGATGGCGATTGCTTTCGTCTGTGGATTAGCGATCCGACCGATTCCTACAGCTGGGCAGGCGTGCTGTGGCGTCCCAATCGTAAAGATTATGAAGTGTATCAGGTGGGGGATCGTCCCGTGTGGGACGAAGTGTTAGATGCGTACAGCCGTTGGGTGGGCTGGGGCCAGCCTGGCCGTGACCGTTTTGGCATGACACTGACCCCGCACCATCAGCACATTTGGCTGGATTCTCCCGACCATCCGTTGCCGTGAcctaggTAATT |
